# Synaptic specificity is collectively determined by partner identity, location and activity

**DOI:** 10.1101/697763

**Authors:** Javier Valdes-Aleman, Richard D. Fetter, Emily C. Sales, Chris Q. Doe, Matthias Landgraf, Albert Cardona, Marta Zlatic

## Abstract

Our nervous system is organized into circuits with specifically matched and tuned cell-to-cell connections that are essential for proper function. The mechanisms by which presynaptic axon terminals and postsynaptic dendrites recognize each other and establish the correct number of connections are still incompletely understood. Sperry’s chemoaffinity hypothesis proposes that pre- and postsynaptic partners express specific combinations of molecules that enable them to recognize each other. Alternatively, Peters’ rule proposes that presynaptic axons and postsynaptic dendrites use non-partner-derived global positional cues to independently reach their target area, and once there they randomly connect with any available neuron. These connections can then be further refined by additional mechanisms based on synaptic activity. We used the tractable genetic model system, the *Drosophila* embryo and larva, to test these hypotheses and elucidate the roles of 1) global positional cues, 2) partner-derived cues and 3) synaptic activity in the establishment of selective connections in the developing nerve cord. We altered the position or activity of presynaptic partners and analyzed the effect of these manipulations on the number of synapses with specific postsynaptic partners, strength of functional connections, and behavior controlled by these neurons. For this purpose, we combined developmental live imaging, electron microscopy reconstruction of circuits, functional imaging of neuronal activity, and behavioral experiments in wildtype and experimental animals. We found that postsynaptic dendrites are able to find, recognize, and connect to their presynaptic partners even when these have been shifted to ectopic locations through the overexpression of receptors for midline guidance cues. This suggests that neurons use partner-derived cues that allow them to identify and connect to each other. However, while partner-derived cues are sufficient for recognition between specific partners and establishment of connections;; without orderly positioning of axon terminals by positional cues and without synaptic activity during embryonic development, the numbers of functional connections are altered with significant consequences for behavior. Thus, multiple mechanisms including global positional cues, partner-derived cues, and synaptic activity contribute to proper circuit assembly in the developing *Drosophila* nerve cord.

## Introduction

Our nervous system is organized into circuits with specifically matched and tuned cell-to-cell connections that are essential for proper function. During development, neurons navigate through the nervous system to reach their target location (Araújo and Tear, 2003;; Dickson, 2002;; Kolodkin and Tessier-Lavigne, 2011;; Tessier-Lavigne and Goodman, 1996;; Yogev and Shen, 2014). Surrounded by numerous cells along their trajectory and in their target area, developing neurons ignore most cells and connect only to specific partners. Furthermore, recent comprehensive electron microscopy (EM) reconstructions of the same circuit in multiple individuals have shown that even the relative numbers of synapses that neurons make with specific partners are very precisely regulated (Eichler et al., 2017;; Gerhard et al., 2017;; Jovanic et al., 2016;; Ohyama et al., 2015). Thus, a neuron can reliably receive 20% of its input from one specific partner, and 5% of input from another, and these fractions of input remain constant across individuals and across different life stages of the same individual. However, we still have an incomplete understanding of the way in which such remarkable synaptic specificity is achieved.

The “lock-and-key” chemoaffinity hypothesis proposes that pre- and postsynaptic partners express specific matching combinations of cell surface molecules that enable them to seek out and recognize each other during development (Langley, 1895;; Sperry, 1963). However, relatively few examples of partner-recognition molecules have been identified to date, so it is an open question whether the use of partner-recognition molecules is a general principle, or whether they are used only in some systems (Hong and Luo, 2014;; Hong et al., 2012;; Krishnaswamy et al., 2015;; Sanes and Yamagata, 2009;; Shen et al., 2004;; Ward et al., 2015). It is also not known whether partner-recognition molecules could specify appropriate numbers of synapses between partners, or whether they just instruct two neurons to form synapses but not how many.

Alternative models propose that pre- and postsynaptic partners seek out specific locations in the nervous system, rather than specific partners (analogous to seeking out a specific address, rather than a person), and once in that location they indiscriminately connect to whichever other neurons are also present there (Peters and Feldman, 1976;; Rees et al., 2017). Consistent with this idea, many neurons have been shown to use positional cues, such as gradients of repellents, to select their termination area (Couton et al., 2015;; Fukuhara et al., 2013;; Mauss et al., 2009;; Sürmeli et al., 2011;; Zlatic et al., 2003, 2009). In such a scenario, additional activity-dependent mechanisms could refine those connections to form functional circuits. Indeed, activity has been shown to refine the patterns of connections in many systems, through Hebbian and/or homeostatic plasticity mechanisms (Giachello and Baines, 2015;; Kaneko et al., 2017;; Marder, 2011;; Schulz and Lane, 2017;; Sheng et al., 2018;; Sugie et al., 2018;; Tien and Kerschensteiner, 2018;; Tripodi et al., 2008;; Turrigiano, 2017;; Yuan et al., 2011). Thus, neurons that fire together have been shown to preferentially wire together in many areas of the vertebrate nervous system through positive feedback (Brown et al., 2009). At the same time, homeostatic mechanisms have been shown to restore activity toward a specific set point through negative feedback, imposing competition and preventing runaway excitation or complete silencing of the circuit (Turrigiano, 2017;; Turrigiano and Nelson, 2004). For example, in the *Drosophila* motor system, inhibition of excitatory premotor neurons increases the numbers of presynaptic specializations in these neurons (Tripodi et al., 2008). In many areas of the cortex, sensory deprivation alters the balance of excitation and inhibition – increasing the strength of excitatory connections onto excitatory neurons (synaptic scaling) and decreasing the strength of inhibitory connections onto excitatory neurons (Kilman et al., 2002;; Maffei and Turrigiano, 2008;; Maffei et al., 2004;; Rutherford et al., 1998). However, the extent to which activity modulates synapse numbers onto excitatory or inhibitory neurons, as opposed to only functional connectivity strength, is still an open question.

Thus, despite recent progress, whether the precise fraction of synaptic input that a neuron receives from a specific partner is determined by specific partner-derived cues, positional cues, activity, or by the interplay between all of these mechanisms is still unknown. It is also unknown if the rules that specify the connectivity between different types of neurons are the same for all neuron types. For example, the rules may differ for connectivity between excitatory and inhibitory neurons, or between distinct types of excitatory or inhibitory neurons. Likewise, it is unclear whether the same mechanisms operate across different phyla. So far, many more examples of activity-dependent connectivity have been observed in vertebrates than in invertebrates.

Finally, a major challenge in the field involves establishing a link between specific defects in structural connectivity, functional connectivity, and behavior. Would developmental defects that result in only minor alterations in fractions of synaptic input from one partner onto another have detectable effects on behavior? In other words, to what extent is it relevant for appropriate function to specify not only who to connect with, but with how many synapses?

These questions have been difficult to address, because studying selective synaptogenesis, including synapse numbers and fractions of total synaptic input, requires comprehensive visualization of synaptic contacts between all of the uniquely identified pre- and postsynaptic partners within a circuit module. We also need to be able to manipulate the activity or location of specific neurons and monitor the effects on structural connectivity, including synapse numbers, within the entire circuit module following such manipulations. Finally, we also need to be able to relate observed structural changes in connectivity to changes in functional connectivity and behavior. However, comprehensive EM reconstruction of entire circuit modules, in wild-type and experimental animals has been out of reach. In addition, genetic tools that allow selective manipulation of uniquely identified circuit elements, and visualization or recording from others were lacking.

To overcome these obstacles, we use the tractable model system, the *Drosophila* larva, where genetic tools for selective manipulation and/or visualization of a large fraction of uniquely identified neurons have recently been generated (Jenett et al., 2012;; Pfeiffer et al., 2008, 2010;; Venken et al., 2011). Furthermore, the larva has a relatively small and compact nervous system and, thanks to recent advances in EM, it has become possible to rapidly image entire regions of its nervous system with synaptic resolution, and to do so in multiple individuals (Gerhard et al., 2017;; Jovanic et al., 2016;; Ohyama et al., 2015). Finally, the larva has a rich behavioral repertoire with well-established quantitative behavioral assays for detecting subtle changes in behavior following various manipulations (Ohyama et al., 2013;; Vogelstein et al., 2014).

We investigated the principles underlying the formation of selective synaptic connections in the somatosensory circuitry in the *Drosophila* larval nerve cord. Recently, we generated comprehensive synaptic-resolution connectivity maps of the circuitry downstream of the mechanosensory Chordotonal (mechano-ch) and nociceptive Multidendritic Class IV (MD IV) neurons in two different individuals (**Figure 1**), which revealed remarkable synaptic selectivity for the strongly connected partners (Jovanic et al., 2016;; Ohyama et al., 2015). Homologous neurons reproducibly make synaptic connections to homologous partners on the left and right sides of the same animal, and in different animals. The fraction of synaptic input a postsynaptic partner receives from a specific presynaptic partner is also conserved across individuals and even across life stages (Gerhard et al., 2017). Previous studies have shown that sensory neurons in this system use positional cues (repellent gradients) to select a specific medio-lateral and dorso-ventral location in the nerve cord where they terminate and from synapses, rather than partner-derived cues (Zlatic et al., 2003, 2009). Thus, selective down-regulation of Robo, the receptor for the glia-derived midline repellent, causes Chordotonal axons to overshoot their usual intermediate target area and terminate more medially. Similarly, overexpression of receptors for the repellent causes Chordotonal neurons to terminate laterally, before they reach their usual target area. However, whether partner-derived cues also exist in this system that would enable the postsynaptic partners to follow their shifted presynaptic partners and connect to them in the ectopic locations was not known. Likewise, the role of activity in circuit assembly in this system was not known. We therefore selectively altered the location or activity of the Chordotonal neurons and generated new EM volumes of the manipulated samples to investigate alterations in connectivity. We complemented these studies by analyzing functional connectivity and behavior following the same manipulations. We show that appropriate postsynaptic cells are able to follow, find, and connect to their displaced presynaptic partners. Thus, when positional cues are altered, partner-derived cues can compensate for their absence and still guide partner recognition. However, displacing Chordotonal neurons led to an alteration in the fractions of input from presynaptic neurons onto postsynaptic neurons. These quantitative defects in connectivity lead to detectable defects in responses to mechanosensory stimuli. Similarly, when Chordotonal neurons were silenced during development, they still formed connections with appropriate postsynaptic partners. However, the fractions of input from presynaptic neurons onto postsynaptic neurons were drastically altered, resulting in significant alterations in functional connectivity within the circuit and behavior. This indicates developing neurons must integrate predefined self-location, partner-derived cues, and partner activity to achieve normal connectivity and function.

**Figure 1.**
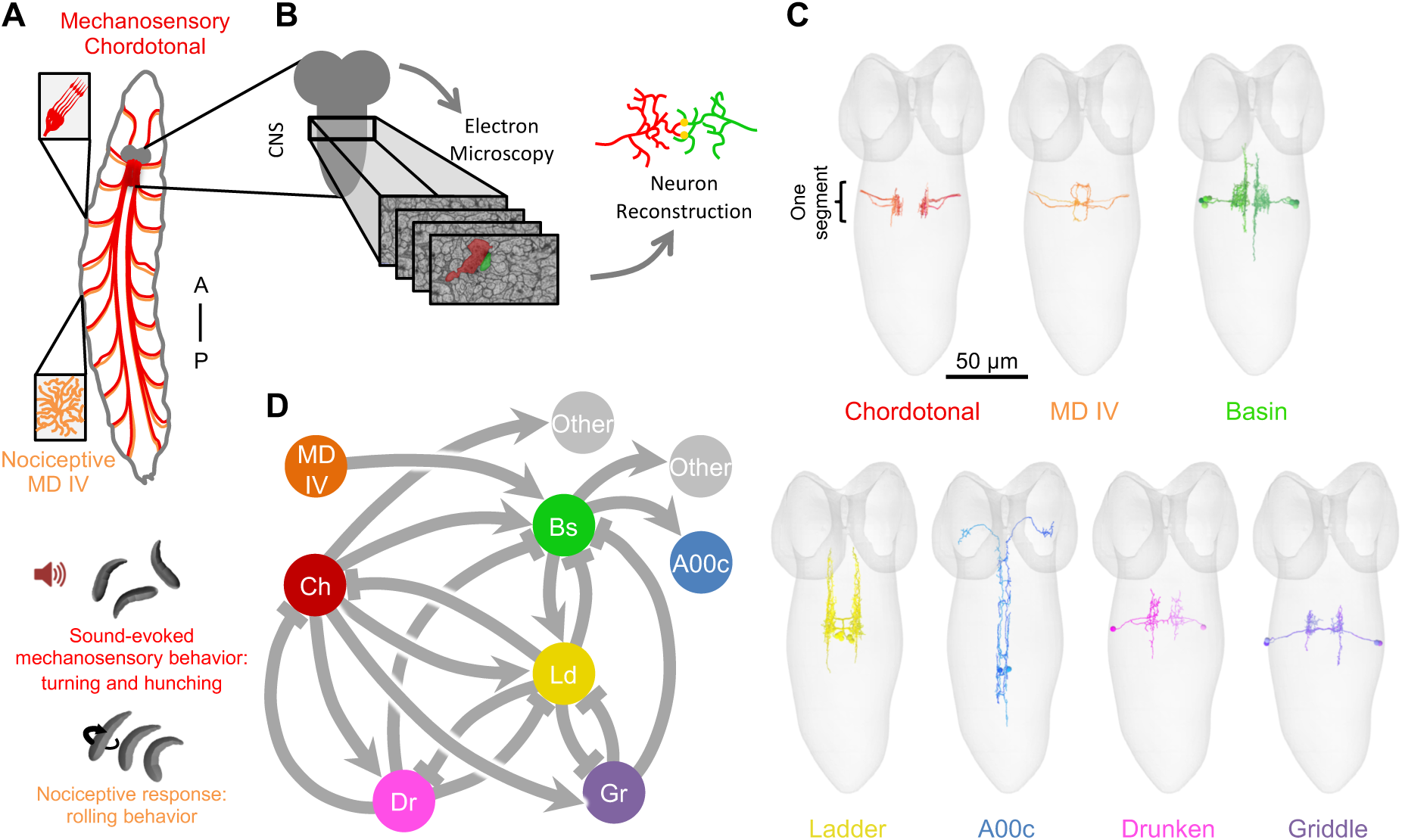
A mechanosensory circuit in *Drosophila* larva revealed by electron microscopy reconstruction. **A)** Schematic of the mechanosensory Chordotonal neurons (red) and the nociceptive multidendritic class IV (MD IV) neurons (orange) spanning the larval body wall and projecting their axons into the CNS. Insets illustrate the morphology of these neurons at the body wall. Vibration generated by sound activates the mechanosensory neurons and elicits a stereotypic behavior consisting of bends and hunches. While activation of MD IV elicits a rolling escape response. **B)** Electron micrographs of thin sections of the CNS allow for high-resolution reconstruction of neurons revealing fine morphology and connectivity. **C)** Skeletonized reconstructions of neurons involved in the mechanosensory circuit generated from EM. Neurons from one abdominal segment (segment A4 for A00c and A1 for all other neurons) of a first instar larva are shown inside the outline of the CNS (gray). Only neurons originating within one abdominal segment are shown for illustration purposes; however, these neurons repeat across multiple segments. These images show 16 individual Chordotonal axons, 6 MD IV axons, 8 Basins, 6 Ladders, 2 A00c, 2 Drunkens and 4 Griddles. **D)** Connectivity diagram of key neurons of the mechanosensory circuit revealed by EM reconstruction. The Chordotonal neurons (red) are direct upstream partners of three groups of inhibitory interneurons: Griddles (purple), Drunkens (pink) and Ladders (yellow). The excitatory Basins cells (green) are a point of multisensory convergence, being directly downstream of the mechanosensory Chordotonals and the nociceptive MD IV neurons (orange). A00c neurons (purple) are excitatory ascending neurons that collect Basin input along the nerve cord and project their axons to the brain. Other neurons downstream of the mechanosensory circuit represent an alternative pathway (gray) not explored in this study. This diagram includes strong synaptic partners that are key for this study, but it does not include all partners. Each circle represents a group of neurons, as opposed to individual neurons. The arrows indicate the direction of the connections: regular arrows represent excitatory connections, and flathead arrows represent inhibitory ones.

## Results

### Postsynaptic partner performs extensive exploration during development

In the embryonic *Drosophila* ventral nerve cord (VNC), somatosensory axons were shown to use positional guidance cues to select where to terminate, branch, and establish synaptic connections (Zlatic et al., 2003, 2009). However, whether their partner dendrites explore their environment seeking out specific presynaptic axons, or whether they connect with whichever axon terminal they contact is unknown. To address this question, we performed live imaging in the intact embryo to follow the development of Chordotonals and one of their preferred postsynaptic partners, Basin neurons (**Figure 2A**).

**Figure 2.**
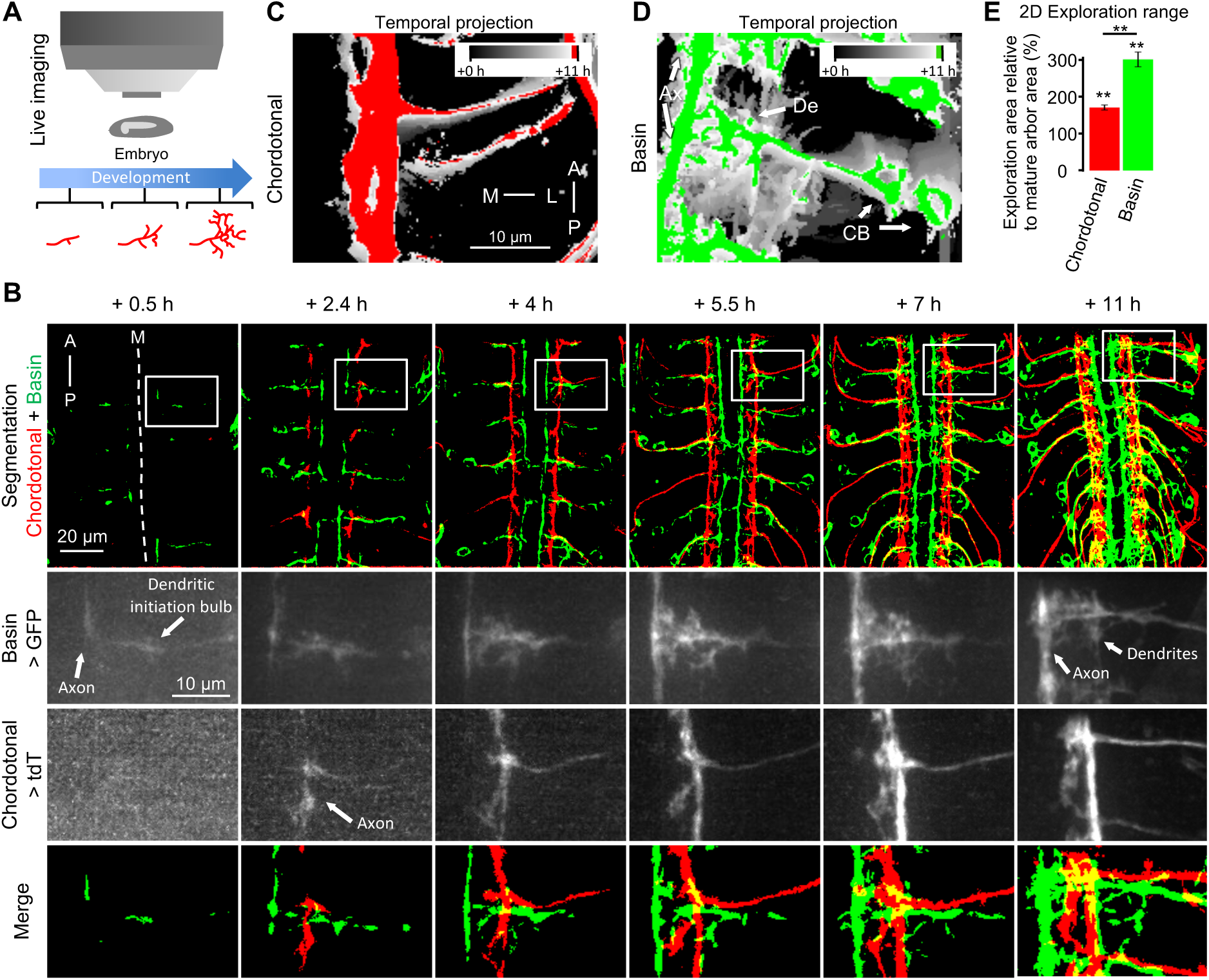
Postsynaptic dendrites have broader exploration range than presynaptic axons. **A)** Schematic of the live imaging experiment. Chordotonal and Basin cells were imaged simultaneously in live embryos throughout development using a spinning-disk confocal microscope. **B)** Developmental time lapse of Chordotonal (red; iav-GAL4 > tdT) and Basin (green; R72F11-LexA > LexAop-GFP) cells. The images are confocal Z-projections of a ventral view of the VNC of live embryos. Time points are relative to the start of the imaging session due to the difference in temperatures before (25 °C) and during imaging (23 °C). The imaging session started 13 hours AEL (time point + 0 h), which is around the moment of earliest detectable expression of GFP in Basin cells. The earliest Basin morphology shows the main branch projecting medially from the cell body to the anteroposterior tract where the axon will form. The swelling in between the Basin cell body and the axon is where the dendrites will branch from. The earliest expression of tdT in the Chordotonal axons was detected around 1 hour after the start of the imaging session. At this moment, the immature Chordotonal axons are already located in the anteroposterior tract where they will span. First row shows several segments in the imaging field of view, subsequent rows show the respective sub regions marked with a white square. Note that the central nervous system normally contracts during development, gradually shifting anteriorly. Due to stochasticity in the driver lines, not all neurons are labelled in all segments. The segmented versions of the light images are included for visual aid, in which Basins are shown in green and Chordotonals in red. Dashed line represents the midline (M). **C)** Temporal projection of Chordotonal axons in one hemisegment during embryonic development. The cumulative Chordotonal exploration area (white) is limited almost exclusively to the area where the mature axons (red) will be present. **D)** Temporal projection of two Basin cells in one hemisegment during embryonic development. The dendrites of Basin cells explore (white) extensively along their hemisegment. They project exploratory filopodia to cover almost their entire hemisegment, sometimes reaching the exploration zone of Basins in the next segment. Basin axons explore much less compared to their dendrites. The mature Basins cease exploring to adopt a final morphology (green) that is much more compact than the cumulative exploration area. CB, cell body; De, dendrites; Ax, axon. **E)** Cumulative two-dimensional exploration area covered during development relative to the area occupied by the mature arbors. Basin dendrites and Chordotonal axons covered a wider cumulative area during developmental exploration than the area occupied by their respective mature arbor (stars above each bar; one-sample t-test with default value of 100%). However, the relative exploration range of Basin dendrites is bigger than that of Chordotonal axons (Wilcoxon test). n= 10 hemisegments each.

At the earliest time we detected fluorophore expression in Basin cells (around 13 hours after egg laying (AEL)), they display a very immature morphology consisting mainly of a bare primary branch projecting from the cell body toward the midline (**Figure 2B**). Approximately in the middle section of this primary branch, there is a swelling which will become the site for dendrite initiation. The leading end at the most medial side of the cell is the growth cone of the Basin developing axon. The cell then proceeds to extend multiple exploratory filopodia from both, its axon and dendrites. These filopodia are short-lived and normally retract within a few minutes, increasing in number and length as development progresses. Dendritic filopodia soon explore most of the anteroposterior length of their hemisegment, occasionally overlapping with other Basin dendrites from neighboring segments. Toward the end of embryonic development (approx. 11 hours after the start of the imaging session), the filopodia exploration ceases and Basins adopt their mature morphology.

At the earliest moment of fluorophore detection in Chordotonal axons, they had already reached their target anteroposterior tract, where they normally arborize (**Figure 2B**). These axons then proceed to extend exploratory filopodia in their immediate vicinity, as they project anteriorly and posteriorly. Interestingly, early Chordotonal axons target the correct anteroposterior tract even before Basin dendritic filopodia initiate exploration, supporting the idea that this axonal targeting is independent of postsynaptic partners (Zlatic et al., 2003, 2009).

Both, presynaptic axons and postsynaptic dendrites display exploratory filopodia during embryonic development. However, the relative two-dimensional (mediolateral and anteroposterior axes) exploration area covered by Basin dendritic filopodia is notably larger than the area covered by Chordotonal axons (**Figure 2C-E**).

Throughout development, dendric and axonal filopodia cover a larger cumulative exploratory area than the final area occupied by their mature arbors (**Figure 2E**). This means many filopodia were not stabilized and covered a space that did not contribute to the final mature morphology. Interestingly, Basin dendritic filopodia repeatedly extended to areas that would eventually not be covered by the mature neuron, presumably areas where partners neurons were absent (**Figure 2D**). This suggests that the dendritic exploration coverage might be broad in nature, and independent of the precise location of presynaptic partners.

In contrast to dendritic exploration, the area covered by Chordotonal axon filopodia during development is almost the same as the area occupied by their mature versions (**Figure 2C**). While this is drastically different to what was observed in the dendritic exploration of Basin cells, it is similar to the exploration of the Basin axons (**Figure 2D**). These results suggest that postsynaptic dendrites might play a more exhaustive role in the exploration for the appropriate presynaptic partners. This is consistent with previous findings that show branching and termination of sensory axons are regulated by the expression of receptors for positional guidance cues independently of their partners (Zlatic et al., 2003, 2009). The Chordotonal axons, whose location is regulated by the global positional cues, appear to provide the instructive signal for postsynaptic dendrites to stabilize those filopodia that contact them.

### Postsynaptic dendrites use partner-derived cues to find and connect to displaced Chordotonal axons

In order to form connections, the presynaptic axon terminals and the postsynaptic dendrites of partner neurons must be in the same location and establish physical contact. In principle, guiding presynaptic axons and postsynaptic dendrites to a precise common location through global positional cues could be sufficient to establish specific connections, making connectivity a result of partner locational coincidence.

However, detailed reconstructions of neuronal maps have provided evidence of striking synaptic specificity (Eichler et al., 2017; Gerhard et al., 2017; Helmstaedter et al., 2013; Jovanic et al., 2016; Kasthuri et al., 2015; Lee et al., 2016; Ohyama et al., 2015; Schneider-Mizell et al., 2016; Takemura et al., 2013, 2015; White et al., 1986; Zheng et al., 2018) that suggests neurons are capable of discriminating the available cells and connecting only with a subset of them. These studies support a more selective mechanism like Sperry’s chemoaffinity hypothesis, which claims there are partner-derived molecular cues that guide the establishment of connections (Sperry, 1963). Thus, positional cues could guide partner neurons to branch in a specific location and then proceed to selectively seek out their partners by searching for specific partner-derived cues.

In order to test the roles of positional guidance cues and putative partner-derived cues in the establishment of synaptic specificity, we genetically induced a shift in location of Chordotonal axons and asked whether their connections with postsynaptic partners would remain intact. We induced this positional displacement by overexpressing the chimeric receptor FraRobo (Bashaw and Goodman, 1999) exclusively in the Chordotonal neurons (**Figure 3A**). This increased their sensitivity to netrin, a midline positional cue, shifting the Chordotonal axons laterally, away from the midline (**Figure 3B**). If neurons search for their specific partners using partner-derived cues, we reasoned that postsynaptic dendrites would find the shifted Chordotonal axons, regardless of their new location, and as long as Chordotonals are still within the range of exploratory dendritic filopodia.

**Figure 3.**
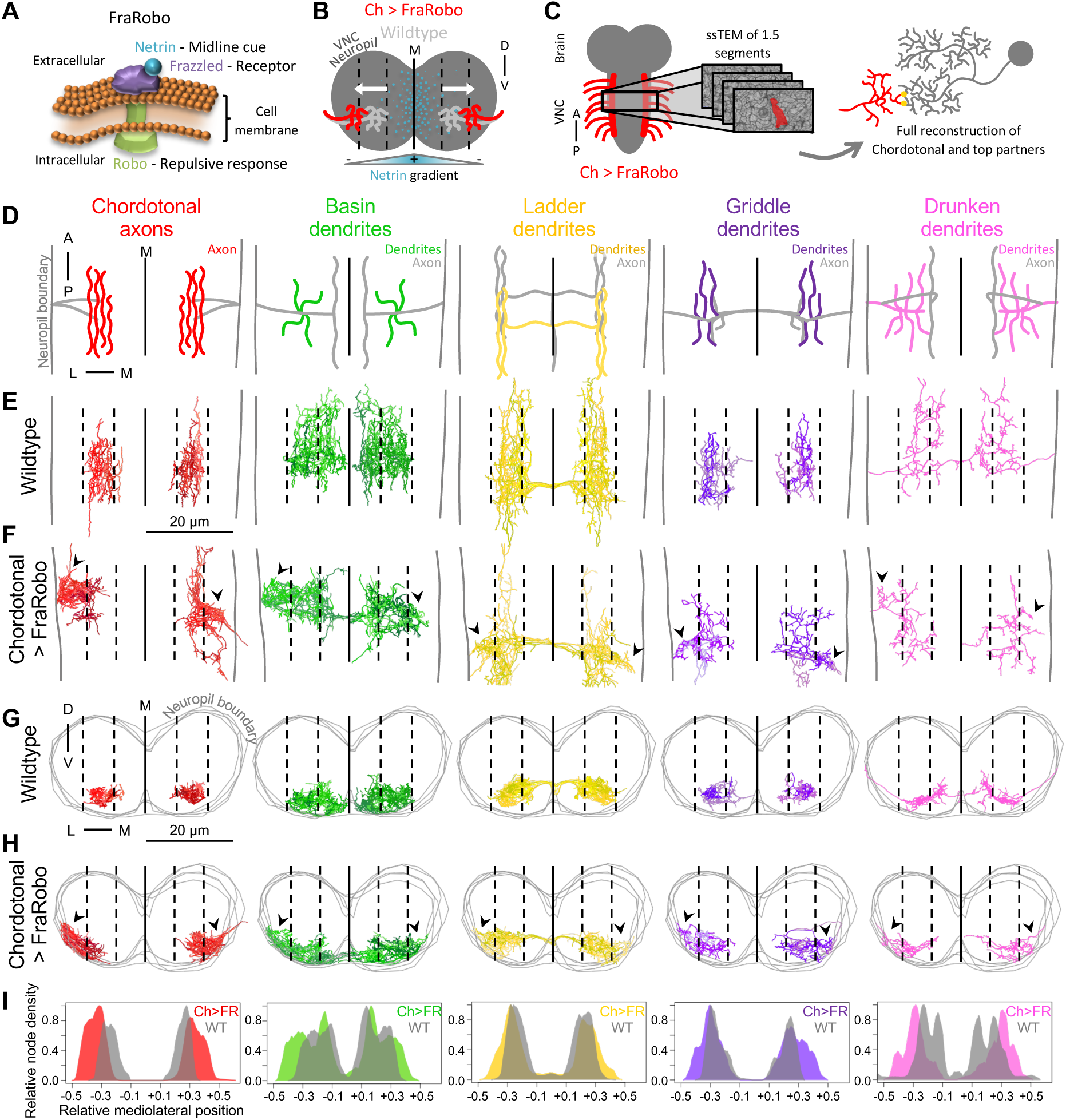
Postsynaptic dendrites extend ectopic branches to reach for the displaced Chordotonal axons. **A)** FraRobo is a chimeric receptor with the ectodomain of Frazzled (purple) and the intracellular domain of Robo (green). The Frazzled component binds to Netrin (blue) while the Robo fraction triggers a repulsive response. **B)** Netrin (blue) is secreted at the midline of the neuropil (dark gray), creating a concentration gradient in the mediolateral axis with the highest concentration at the midline (M) and the lowest at the lateral ends. The Chordotonal (Ch) axons that ectopically express FraRobo (red) are more sensitive to Netrin and avoid areas of high concentration of it, resulting in a lateral shift compared to wildtype Chordotonal axons (light gray). **C)** 1.5 segments of the VNC of an animal expressing FraRobo exclusively in the Chordotonal neurons was imaged using serial section transmission electron microscopy (ssTEM). Chordotonal neurons and their top partners were fully reconstructed to investigate the possible effects of this manipulation on morphology and connectivity. **D)** Schematic of a dorsal view of the Chordotonal axons and the dendritic regions of their postsynaptic partners in one abdominal segment. The colored subcellular regions correspond to the subarbors displayed in subsequent panels in this figure, those regions in gray are not shown. **E-H)** Dorsal (E and F) and cross section (G and H) views of the reconstructed Chordotonal axons and postsynaptic partner dendrites in wildtype (E and G) and in a sample in which Chordotonals express FraRobo (F and H) (*Ch-GAL4 > UAS-FraRobo*). Chordotonal axons expressing FraRobo are displaced laterally (arrowheads), away from the midline (M; solid line), reaching the edge of the neuropil. The postsynaptic partners display ectopic branches in lateral domains (arrowheads) as a consequence of the displacement of their presynaptic partner. The neuropil boundary is represented by either a pair of gray vertical lines for dorsal views (D-F) or gray consecutive rings for cross section views (G and H). Dashed lines split the maximum width of the neuropil in six equidistant sections, three on either side of the midline. **I)** Node density distributions of reconstructed neurons in the mediolateral axis. The node density of all the reconstructed neurons was quantified using a 2.5 μm sliding window across the mediolateral axis. The mediolateral positions are normalized to the width of the neuropil of the corresponding EM volume. The midline is represented by zero in the X-axis. The node densities are normalized to the maximum density of the respective cell type.

The best way to confidently determine if the modified location of Chordotonal axons affects their connectivity is to look for any morphological changes in their synaptic partners and quantify the connections with them. In order to do this, we performed EM reconstruction of the FraRobo-expressing Chordotonal neurons and their key postsynaptic partners in a volume spanning 1.5 abdominal segments of the VNC of a first instar larva (**Figure 3C**). EM reconstructions revealed that the Chordotonal axons expressing FraRobo were indeed shifted laterally closer to the boundary of the neuropil (**Figure 3D-H**). These shifted axons grouped together in semi-isolated lateral clusters, losing their anteroposterior continuity that is normally observed between segments (**Figure 3E, G**). The expression of FraRobo affected different Chordotonal axons with different magnitudes, causing some Chordotonals to shift more than others. This resulted in some axons not shifting or occupying both their normal location and the shifted one.

Surprisingly, the displacement of the Chordotonal axons caused a subsequent lateral shift of the dendrites of the postsynaptic partners. The excitatory Basin neurons normally receive Chordotonal input in the medial and lateral subregions of their dendritic arbors. When the Chordotonal axons were shifted laterally, Basins broadened their dendritic coverage by spreading out their dendrites laterally to reach for the ectopic Chordotonal input (**Figure 3E-H**). Basin dendrites were now found all the way to the edge of the neuropil, a location where they are normally absent. Ladders, Griddles and Drunkens are inhibitory interneurons, predominately downstream of Chordotonal neurons. When the Chordotonal axons were shifted laterally, these inhibitory interneurons also extended discrete ectopic dendrites reaching for the displaced Chordotonal input (**Figure 3F, H**).

In fact, we were able to reproduce an analogous postsynaptic dendritic displacement as a consequence of presynaptic shift in a different and completely independent pair of synaptic partners in the VNC: presynaptic dbd sensory neurons and postsynaptic A08a interneurons. Sales and colleagues have shown that selectively expressing Robo-2 or Unc-5 in dbd neurons caused an intermediate or strong lateral shift of their axons, respectively (**Figure 4A-C**) (Sales et al., 2019). Interestingly, we also observed a subsequent lateral shift in the dendritic distribution of A08a as a consequence of the lateral displacement of the presynaptic dbd neurons (**Figure 4A-E**). This is consistent with the extension of ectopic dendritic branches from Basin, Ladder, Drunken and Griddle, when the presynaptic Chordotonal axons were shifted laterally. We were able to reproducibly detect this displacement across multiple light-level microscopy samples of A08a and Basin dendrites (**Figure 4E, F**). This striking morphological adaptation of the postsynaptic interneuron dendrites in response to the displacement of the presynaptic Chordotonal or dbd axons indicates these interneurons must use partner-derived cues that guide them to their partners at their new unusual location.

**Figure 4.**
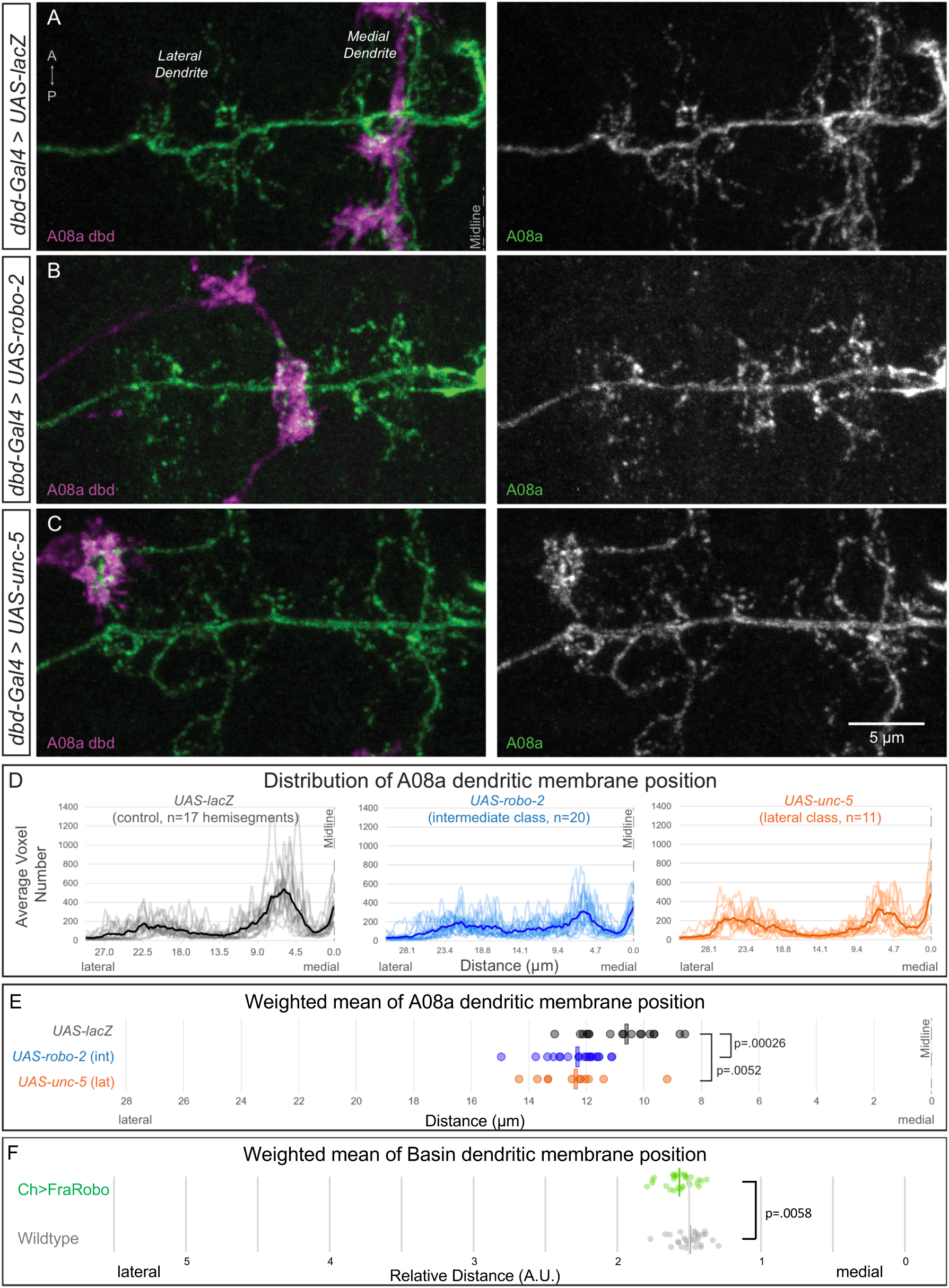
Displacement of two different types of sensory neurons cause their respective postsynaptic partners to shift the distribution of their dendrites. **A-C)** Confocal maximum intensity projection of the dorsal view of dbd axon terminal (magenta) and the A08a dendritic domain (green) in one hemisegment of a 3^rd^ instar larva. Merged channels shown to the left; A08a channel shown to the right. **A)** In wild type, dbd axon targets the A08a medial arbor. n=17 hemisegments from 11 animals. **B)** In animals in which dbd express Robo-2, dbd axons are shifted laterally and often contact the A08a intermediate domain. n=20 hemisegments from 10 animals. **C)** In animals in which dbd express Unc-5, dbd often contacts the A08a lateral arbor. n=11 hemisegments from 10 animals. **D)** Quantification of the distribution of A08a dendrite position in the context of medial, intermediate, and lateral innervating dbd neurons. Each transparent line represents one hemisegment and each solid line represents the average for the cohort. **E)** Weighted mean of the dendrite distributions shown in D. Each circle represents the weighted mean for each hemisegment and the bar represents the average weighted mean for each cohort. Average weighted mean: *UAS-lacZ*: 10.63 µm; *UAS-robo-2*, intermediate: 12.33 µm; *UAS-unc-5*, lateral: 12.38 µm. P-values were obtained using an unpaired t-test. **F)** Weighted mean of the distribution of Basin dendrites in animals in which Chordotonals express FraRobo (*Ch-GAL4>UAS-FraRobo, UAS-tdT, Basin-LexA>LexAop-GFP*) and control (*Ch-GAL4>UAS-tdT, Basin-LexA>LexAop-GFP*). The mediolateral axis was normalized by the distance between Basin axons from left and right sides to correct for slight differences in size of sample stretching. n= 30 hemisegments each.

### Interneurons axons and dendrites independently reach for displaced Chordotonal axons

Since most of the Chordotonal synapses onto their postsynaptic partners are located in the dendrites (axo-dendritic connections), it is unclear whether the interneuron axons would also be shifted, as were the dendrites when Chordotonals expressed FraRobo. Additionally, previous work shows that sensory axons can be displaced regardless of their postsynaptic partner’s location (Zlatic et al., 2003, 2009). This raises the possibility that interneuron axons and dendrites might use different guidelines when they are guided to a specific location during development.

To investigate this further, we looked at the location of the axons of the neurons whose dendrites are postsynaptic to the shifted Chordotonals (**Figure 5A**). Interestingly, the lateral shift of the Chordotonal axons caused a subsequent shift in some, but not all, of the axons of key downstream partners. The axons of Ladder and Drunken showed clear ectopic branches that overlap with the shifted Chordotonal axons (**Figure 5C** and **Figure 5E**). Interestingly, the ectopic axonal branches of Ladder and Drunken are also presynaptic to Chordotonals (forming axo-axonic connections) and some of the other shifted interneuron dendrites (axo-dendritic connections) (**Figure 5G**). This suggests the interneuron axons were shifted to form ectopic synapses to/from Chordotonals, interneurons or both. Interestingly, a portion of Ladder axons in one hemisegment was not shifted (upper right, **Figure 5C**), corresponding to the same location where Chordotonal axons were least shifted (**Figure 3F**). While this variability was expected, it served as an internal (same sample) control of the downstream effects of this manipulation. This shows that the formation of ectopic axons in Ladder neurons is a direct result of the lateral displacement of their main presynaptic partner, the Chordotonal neurons.

**Figure 5.**
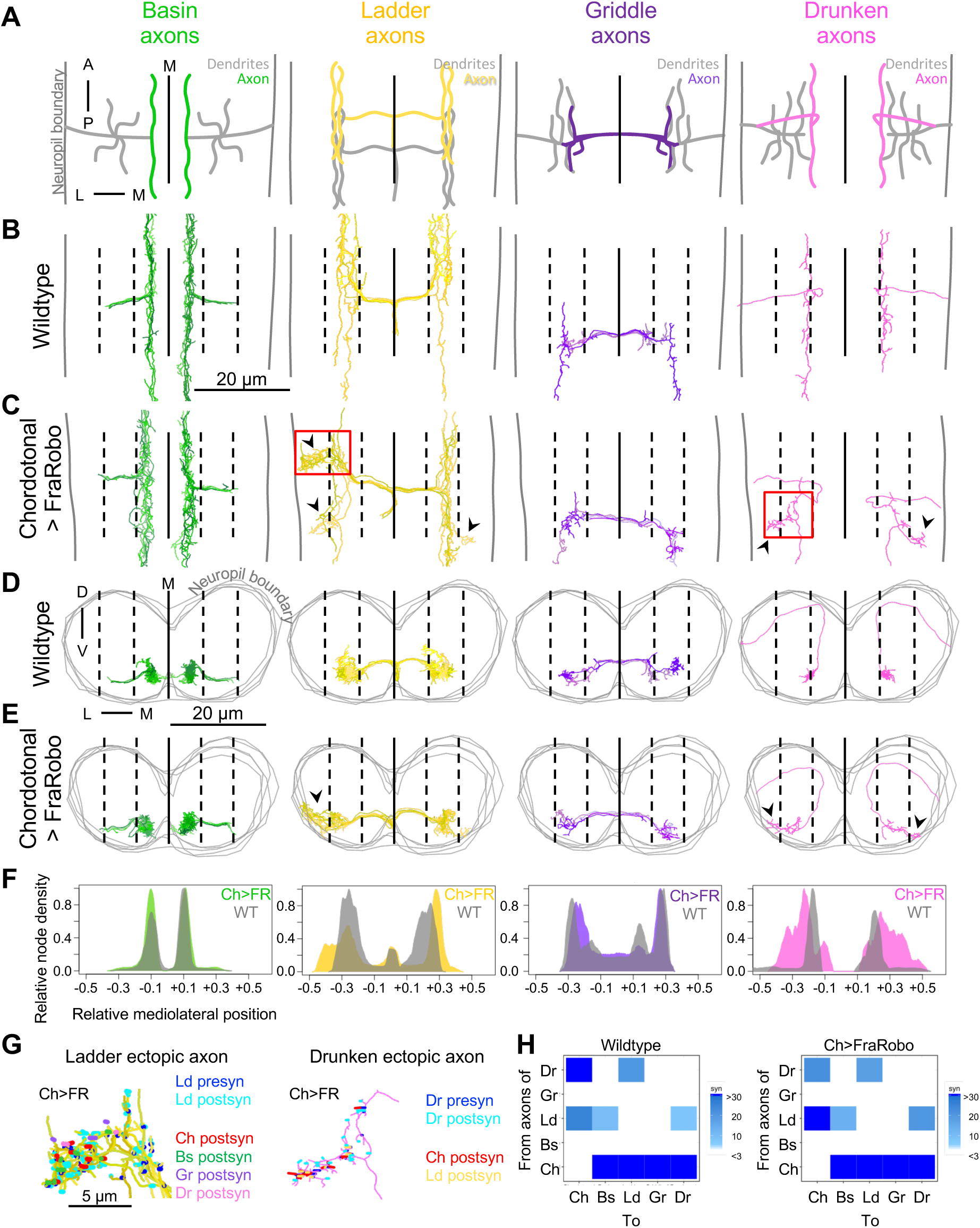
Interneuron axons reach for displaced Chordotonal axons, independently of their dendrites. **A)** Schematic of a dorsal view of the axonal regions of the Chordotonal postsynaptic partners in one abdominal segment. The colored subcellular regions are consistent with those displayed in subsequent panels in this figure. **B-E)** Dorsal (B and C) and cross section (D and E) views of the reconstructed axons of interneurons (Chordotonal partners) in wildtype (B and D) (*w1118*) and in a sample in which Chordotonals express FraRobo (C and E) (*Ch-GAL4 > UAS-FraRobo*). The axons of Ladder and Drunken extend ectopic branches (arrowheads) due to the displacement of the Chordotonal axons. However, Basin and Griddle axons do not display any lateral displacement. The neuropil boundary is represented by either a pair of gray vertical lines for dorsal views (B and C) or gray consecutive rings for cross section views (D and E). Dashed lines split the maximum width of the neuropil in six equidistant sections, three on either side of the midline (M; solid line). **F)** Node density distribution of reconstructed axons in the mediolateral axis. The node density of all the reconstructed neurons was quantified using a 2.5 μm sliding window across the mediolateral axis. The mediolateral positions are normalized to the width of the neuropil of the corresponding EM volume. The midline is represented by zero in the X-axis. The node densities are normalized to the maximum density of the respective cell type. **G)** Ectopic axonal branches of Ladder and Drunken from C (red squares) with synapses color-coded by partner, showing they are presynaptic to shifted Chordotonal axons and to other Chordotonal partners. This means that the axonal shift of Ladder and Drunken could be directly caused by the shifted Chordotonal axons and/or by the indirectly-shifted interneurons. **H)** Connectivity matrix of axon-to-whole-neuron connections between Chordotonal (Ch), Basin (Bs), Ladder (Ld), Griddle (Gr) and Drunken (Dr). Axons of Basin and Griddle do not normally synapse onto each other or any neuron that was displaced as a consequence of the expression of FraRobo in Chordotonals (i.e. Chordotonal, Ladder or Drunken). Matrices only include connections between cell types with 3 or more synapses to a single neuron.

Contrastingly, the axons of Basins and Griddles were not displaced (**Figure 5C, E**). Since Basin and Griddle do not normally form synapses with Chordotonals or any of the other shifted interneurons (Basin, Ladder, Griddle or Drunken) (**Figure 5H**), their axons did not need to connect to any ectopic partner, explaining why the location of their axons was not affected. The intact locations of these axons serve as a control to rule out any major developmental defect that could have generated an overall shift in the entire neuropil. These results indicate that, similarly to dendrites, interneuron axons also use partner-derived cues to find their partners regardless of their precise location. However, the partner-derived cues interneuron dendrites follow are independent of those used by the axon of the same cell.

### Shifted Chordotonals retain strongly connected partners and do not gain new partners

EM reconstruction showed that the shift of Chordotonal axons caused a subsequent shift in their postsynaptic partners. However, we then asked whether the connectivity between these partner neurons was preserved. We not only looked at the connectivity with key partners, we also reconstructed to identification all neurons downstream of the shifted Chordotonal axons (**Figure 6A**). Some partners neurons could not be identified because they exited the EM volume and the contained fragments were too small to be morphologically matched to a wildtype reference. We found that the shifted Chordotonal neurons remain connected to most of their key partners (**Figure 6B-C**). However, the ranking of the Chordotonal downstream partners was significantly redistributed. For example, in wildtype, Ladder and Griddle are the predominant top partners (by total input) downstream of Chordotonals. Interestingly, when Chordotonals are shifted by the expression of FraRobo, Basins become the top partner while Ladders are demoted down the ranking. This redistribution of the connectivity ranking shows that the physical displacement of the presynaptic partners has a significant impact in connectivity.

**Figure 6.**
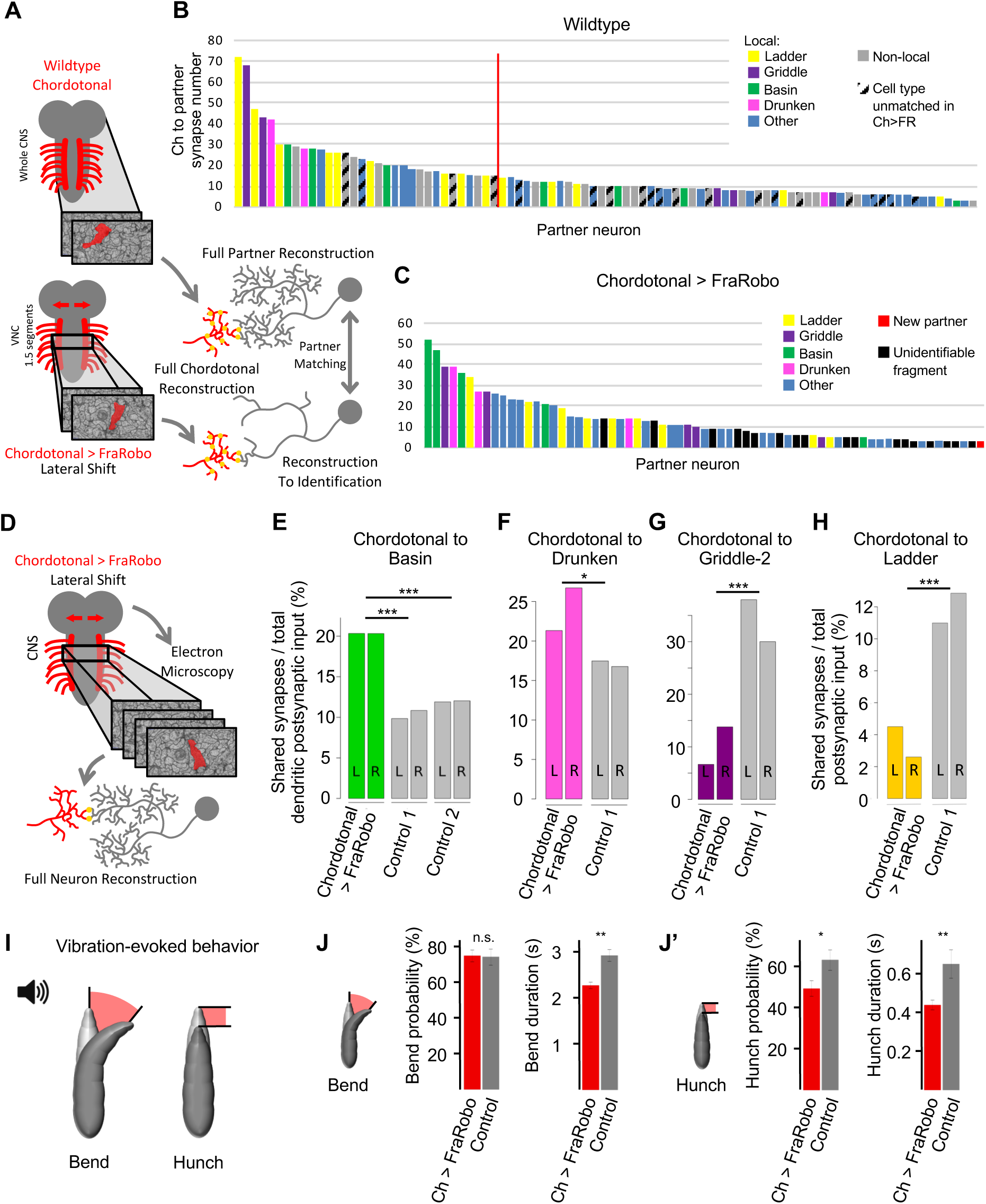
Shifting the location of the Chordotonal axons alters the circuit connectivity, generating deficient mechanosensory behavior. **A)** Chordotonal neurons and downstream partners were reconstructed in two different EM volumes: one wildtype (*w1118*) and one in which the Chordotonals expressed FraRobo (*Ch-GAL4 > UAS-FraRobo*). The wildtype volume encompasses the whole CNS, and most of the partners downstream of Chordotonals in it have been fully reconstructed. The FraRobo volume consists of approximately 1.5 abdominal segments. In this volume, the non-preferred (i.e. not Basin, Ladder, Griddle, and Drunken) Chordotonal partner neurons were partially reconstructed up to a point where they could be identified and matched to their fully reconstructed neurons counterparts from the wildtype volume. **B-C)** Connectivity plots of the downstream partners of Chordotonal neurons from one hemisegment in a wildtype EM volume (B) and a *Ch-GAL4 > UAS-FraRobo* volume (C). Bars represent individual neurons. Number of synapses shown are from all eight grouped Chordotonal axons onto individual postsynaptic partners. **B)** Local neurons were defined as those within the same region (same segment and half of the two adjacent segments) as Chordotonals (limits set to approximate those of the FraRobo volume). Those neurons partially within these limits were considered as local if the encompassed fractions could still be identified, otherwise they were considered as non-local. Those cell types that were not found downstream of Chordotonal neurons in the FraRobo volume are marked as unmatched. Only those neurons with at least 3 synapses from Chordotonals on each side (left and right) of the segment are shown. Red vertical line indicates the threshold between neurons strongly (≥15 total synapses) and weakly (<15 total synapses) connected to Chordotonals. **C)** The ranking of Chordotonal downstream partners is redistributed compared to wildtype. Most of top local partners (from B) remain connected when Chordotonals were shifted. Partial fragments of neurons that leave the EM volume that could not be identified are colored in black (correspond to “unmatched” in B). Neurons downstream of Chordotonals in the FraRobo volume (reproducible in left and right) that are not downstream Chordotonals in the wildtype volume are marked in red. Only those neurons with at least 3 synapses from Chordotonals are shown. **D)** Chordotonals and their key downstream partners (Basin, Ladder, Griddle and Drunken) were fully reconstructed in a *Ch-GAL4 > UAS-FraRobo* EM volume of 1.5 segments. **E-H)** Connectivity between FraRobo-expressing Chordotonal neurons and key postsynaptic partners is altered. The number of synapses from Chordotonals onto the postsynaptic partner was divided by the total number of dendritic inputs of the postsynaptic partner (E-G), or total postsynaptic input (H) (dendritic and axonal input considered for Ladder due to strong Chordotonal input onto both subcellular compartments). Connectivity from neurons in the right (R) and left (L) sides of one segment is shown separately to show consistency within sample. However, synapse counts from right and left were grouped for statistical analysis. Connectivity fractions were compared using Chi-square test. **I)** Sound-generated vibration activates Chordotonal neurons and elicits bending and hunching behaviors. **J-J’)** Animals with shifted Chordotonal axons have deficient mechanosensory behavioral responses. The probability of turning behavior in animals with shifted Chordotonals is similar to that of controls but with longer duration (J). However, the probability and duration of hunch are both increased in animals with shifted Chordotonals (J’). Error bars represent the 95% confidence interval for probabilities or standard error for durations. For probabilities: experimental (red) n= 677; control (gray) n= 367. For duration: experimental (red) n= 506; control (gray) n= 272.

In addition to rearranging the order in the ranking of Chordotonal partners, we asked whether the lateral displacement of Chordotonal axons led to the loss of any postsynaptic partners. In fact, we found a lower number of total postsynaptic partners in the FraRobo EM volume. However, we believe this to be, at least partially, because of a slight age difference between samples, as younger animals have smaller neurons with fewer synapses (Gerhard et al., 2017).

Therefore, Chordotonal neurons in the FraRobo volume have fewer partners, mostly a result of fewer noisy connections to weakly connected neurons. Strikingly, most of strongly connected cell types downstream of Chordotonals in wildtype (**Figure 6B**) remain connected in the FraRobo volume (**Figure 6C**). This indicates strongly connected partners must have arisen from a specific affinity to Chordotonals and not from locational coincidence alone.

Alternatively, we asked whether the lateral shift of Chordotonal axons led to the gain of new synaptic partners found at their new location, as would be expected from a purely location-based mechanism for synaptic specificity. Strikingly, we found only a single cell type downstream of the shifted Chordotonal axons that is not a regular partner in the wildtype (**Figure 6C**). This new partner was weakly connected and barely above the connectivity threshold. This extremely low number of new partners and the retention of strongly connected partners shows remarkable partner specificity despite of the altered location of the Chordotonal neurons.

### Shifting location of Chordotonal axons alters structural connectivity within the circuit

We then asked whether the Chordotonal key postsynaptic partners received a normal amount of Chordotonal input or the connectivity balance between them was lost. Since the total number of synapses in the nervous system increases throughout development of the larva (Gerhard et al., 2017), it is not possible to use this direct measurement to accurately compare connectivity strength across different EM volumes. However, the fraction of input a neuron receives from (or makes onto) a specific partner has been shown to be remarkably conserved across individuals and development (from first to third instar larva in *Drosophila*) (Gerhard et al., 2017). Therefore, we report connectivity between partner neurons (for example: neuron A synapsing onto neuron B) across volumes as fractions, resulting from dividing the number of shared synapses by the total postsynaptic input (synapses from A to B/total incoming synapses of B). We fully reconstructed the preferred Chordotonal partners (Basin, Ladder, Drunken, and Griddle) in the FraRobo volume and compared their fractions of Chordotonal input to those in wildtype (**Figure 6D**).

Basin, Drunken and Griddle receive most of their Chordotonal input onto their dendrites, therefore we calculated the connectivity from Chordotonals relative to the total amount of input onto their dendrites, which are fully contained in the FraRobo EM volume. However, Ladders normally receive a significant amount of Chordotonal input onto both, their dendrites and axons. Therefore, we calculated the fraction of total (axonal and dendritic) input synapses for Ladders. Since parts of Ladder arbors exited the FraRobo EM volume, equivalent (in coverage) subvolume limits were used to restrict the total number of Ladder input synapses considered from the wildtype volume, in order to simulate a subvolume similar to the one for FraRobo. This correction made it possible to compare Ladder connectivity fractions between wildtype and FraRobo volumes. We found the fractions of Basin and Drunken input from Chordotonal neurons were higher than in control volumes (**Figure 6E-F**). Contrastingly, the fractions of Griddle and Ladder input from Chordotonal neurons were lower than control (**Figure 6G-H**). Altogether, we found that the connectivity balance between shifted Chordotonals and key postsynaptic partners was disturbed, as some connections increased, and others decreased, compared to wildtype.

Thus, the EM reconstruction of the partners of Chordotonal neurons revealed that dendrites and axons of interneurons are able to find and connect to the presynaptic Chordotonal axons, even when these have been shifted to ectopic locations. However, the analysis of detailed synaptic connectivity in such animals revealed that the relative Chordotonal input onto different postsynaptic neurons is significantly different to wildtype.

### Shifting location of Chordotonal axons generates deficient mechanosensory behavior

Chordotonal neurons are activated by sound-generated vibration, which elicits stereotypic body-bending and hunching behaviors in larvae (Jovanic et al., 2016; Ohyama et al., 2015). Therefore, we measured these behavioral responses to vibration to test whether the altered connectivity of the shifted Chordotonal circuit has an effect on the overall functional output (**Figure 6I**). Larvae with FraRobo-expressing Chordotonals displayed bending and hunching behavioral responses to vibration, indicating Chordotonals are still functional and sensitive to this stimulus. However, bending responses were significantly shorter in duration compared to controls (**Figure 6J**). Similarly, the probability and duration of hunches were both significantly lower than in controls (**Figure 6J’**). This shows that despite the displacement of the Chordotonal neurons, the mechanosensory circuit is still functional and capable of generating mechanically-evoked behavior. Nevertheless, the overall circuit connectivity was altered by the locational shift of Chordotonal axons, generating deficient mechanosensory behavioral responses. Therefore, despite evident partner specificity, precise positioning of synaptic partners has a significant impact on circuit assembly.

### Lack of Chordotonal activity during development alters connectivity in the circuit

We showed that Basin dendrites project long exploratory filopodia and establish their first contacts with Chordotonal axons during late embryonic development. These interactions begin right before the first action potentials happen in the developing nervous system of the *Drosophila* embryo (Baines and Bate, 1998). This raises the possibility that neuronal activity between developing neurons might contribute to wiring specificity (Akin et al., 2019), in addition to specific partner-derived molecular cues. Despite extensive work investigating the role of activity in circuit formation, it is still unclear whether activity is involved in partner specificity (Kaneko and Ye, 2015; Sugie et al., 2018).

To investigate the role of neural activity during development in the establishment of synaptic specificity, we permanently blocked synaptic transmission in the Chordotonal neurons through the targeted expression of tetanus toxin light chain (TNT) (Sweeney et al., 1995). In order to assess the effect of this manipulation on the circuit’s connectivity, we performed EM reconstruction of silenced Chordotonals and their key downstream partners in an EM volume that spans 1.5 abdominal segments of a first instar larva (**Figure 7A**). We found that silencing the Chordotonal neurons had an effect on the connectivity from Chordotonals onto Basin, Griddle, and Ladder neurons, albeit in opposite ways. The fraction of Basin input synapses from inactive Chordotonals was higher than in controls (**Figure 7B**). In contrast, the fraction of Griddle and Ladder input from Chordotonals was decreased (**Figure 7D-E**), while there was no difference in the connectivity between Chordotonal and Drunken neurons (**Figure 7C**). EM reconstruction revealed that silent Chordotonals and key downstream partners remain connected; however, the relative number of connections between them was different across cell types compared to wildtype.

**Figure 7.**
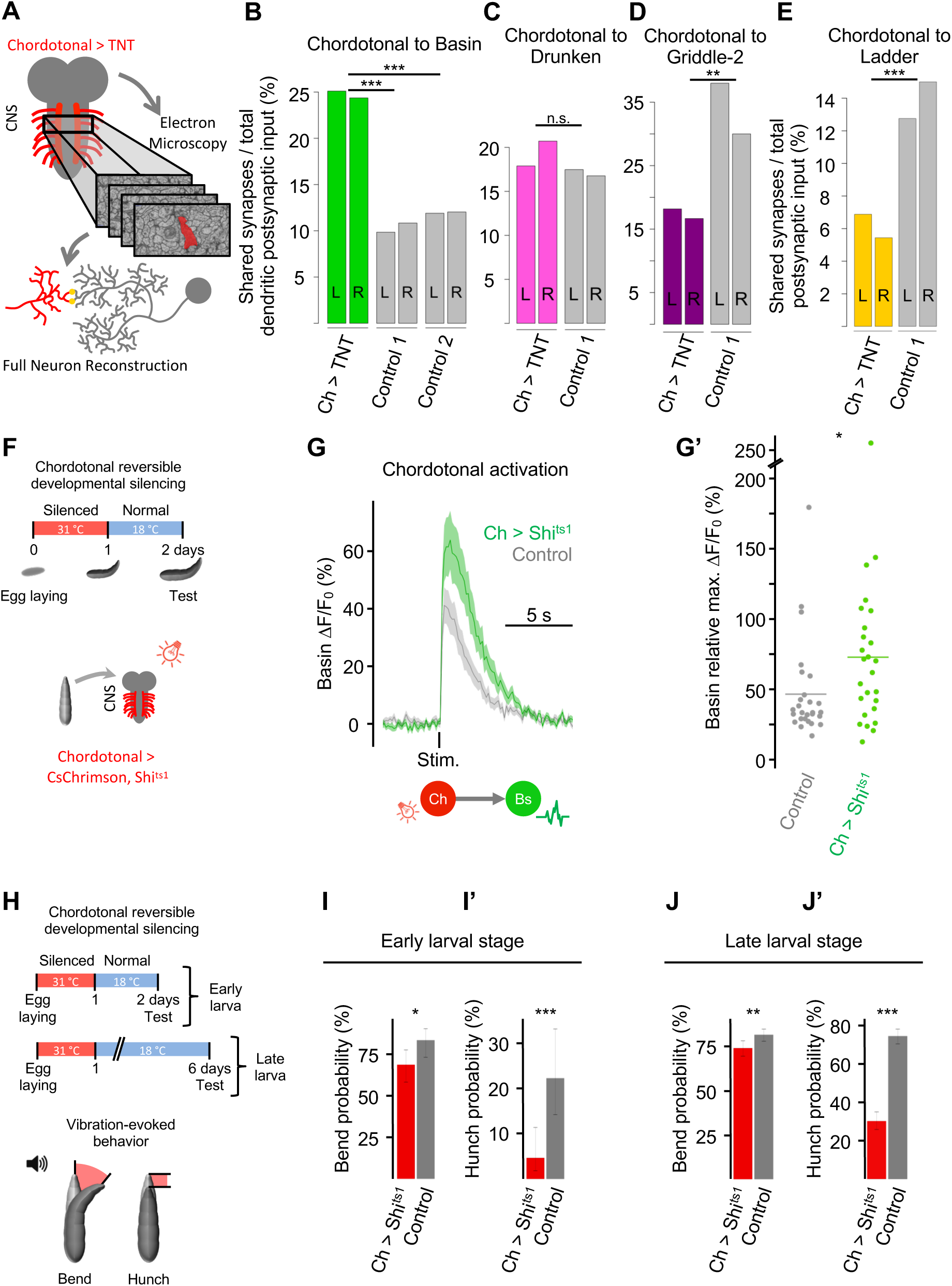
Lack of Chordotonal input during development alters the connectivity of the circuit and generates defective behavior. **A)** Chordotonal neurons and downstream partners were reconstructed in two different EM volumes of first instar larvae: one wildtype (*w1118*) and one in which the Chordotonals were silenced through the expression of TNT (*Ch-GAL4 > UAS-TNT*). The TNT volume consists of approximately 1.5 abdominal segments of the VNC. In this volume, the key (i.e. Basin, Ladder, Griddle, and Drunken) Chordotonal partner neurons were fully reconstructed. **B-E)** Connectivity revealed by EM reconstructions of Chordotonals (Ch) silenced with TNT and key postsynaptic partners in one abdominal segment of a first instar larva. To report the relative number of connections between partner neurons, the number of synapses from Chordotonals onto the postsynaptic partner was divided by the total number of dendritic (B, C, D), or dendritic and axonal (E) inputs of the postsynaptic partner. The experimental animal’s genotype was *UAS-TNT > Ch-GAL4*. Controls were animals of the *w1118* genotype. Connectivity from neurons in the right (R) and left (L) sides is shown separately. However, left and right synapse counts were grouped for statistical analysis using Chi-square test. **F)** Schematic of the experimental conditions for reversible silencing of Chordotonal neurons during development. Embryos with Chordotonal neurons expressing Shi^ts1^ and CsChrimson were incubated at 31 °C (restrictive temperature) for 24 hours and then transferred to 18 °C (permissive temperature) for another day before testing. Isolated CNS were used to record calcium responses in Basins (G-G’) to the optogenetic activation of Chordotonal neurons. **G)** Basin calcium responses (mean ± s.e.m) to Chordotonal optogenetic activation (Stim., 1040 nm for 100 ms) increase when the Chordotonal neurons were reversibly silenced with Shi^ts1^ during development (green trace; *Basin-GAL4 > UAS-GCaMP6s, Ch-LexA > LexAop-CsChrimson, LexAop-Shi^ts1^*) compared to control (gray trace; *Basin-GAL4 > UAS-GCaMP6s, Ch-LexA > LexAop-CsChrimson*). **G’)** Quantification of the calcium responses in G. Relative maximum ΔF/F0 values had the trial-specific baseline subtracted. n= 11 animals for experimental (green); n= 12 animals for control (gray). **H)** Schematic of the experimental conditions for reversible silencing of Chordotonal neurons during development for behavioral experiments. Embryos with Chordotonal neurons expressing Shi^ts1^ were incubated at 31 °C (restrictive temperature) for 24 hours and then transferred to 18 °C for another day (I-I’) or 5 days (J--J’) before testing. Animals were stimulated with sound-generated vibration (1000 Hz tone). **I-J’)** Reversibly silencing Chordotonal cells during development significantly reduces the behavioral responses to vibration of early stage larvae (I-I’), with persisting defects in late stage larvae (J-J’). Larvae in which Chordotonal neurons were silenced during development have lower probability for bending (I, J) and hunching (I’, J’) responses. Experimental (red) genotype: *Ch-GAL4 > UAS-Shi^ts1^*. Control (gray) genotype: *+ > UAS-Shi^ts1^*. Error bars represent the 95% confidence interval. **I-I’)** The probabilities for bending and hunching behaviors are lower in early stage larvae in which the Chordotonal neurons were silenced (red) during development than in controls (gray). n= 86 animals for experimental; n= 72 for control. **J-J’)** The probabilities for bending and hunching behaviors are lower in late stage larvae in which the Chordotonal neurons were silenced (red) during development than in controls (gray). n= 380 animals for experimental; n= 476 for control.

### Reversibly silencing Chordotonal neurons during development alters functional connections with downstream partners

We tested whether the significant differences in structural connectivity induced by silencing Chordotonal neurons also result in differences in functional connectivity. Since the Chordotonal neurons in this EM volume are permanently silenced by the expression of TNT, it is impossible to use this same manipulation to test for functional connectivity with any postsynaptic partner. We therefore genetically targeted the overexpression of temperature-sensitive Shibire (Shi^ts1^) in the Chordotonal neurons to reversibly block synaptic vesicle endocytosis (Kitamoto, 2001). This analogous manipulation allowed us to temporarily block synaptic transmission during embryonic development and later restore activity to test functional connectivity with postsynaptic partners (**Figure 7F**).

We used CsChrimson, a red-shifted channelrhodopsin (Klapoetke et al., 2014), to optogenetically activate Chordotonal neurons. Simultaneously, we expressed GCaMP6s (Chen et al., 2013), a fluorescent calcium indicator, in Basins to monitor their responses to the optogenetic activation of Chordotonals. We found Basin calcium responses to the activation of Chordotonals to be greater when Chordotonals were inactive during development, compared to animals in which Chordotonals activity was never manipulated (**Figure 7G-G’**). This functional connectivity increase is consistent with the observed increase in structural connectivity between TNT-inactivated Chordotonals and Basin neurons revealed by EM reconstructions (**Figure 6E**).

### Lack of Chordotonal activity generates long-lasting deficient mechanosensory behavior

We then investigated whether the significant differences in structural and functional connectivity induced by silencing of Chordotonal neurons also result in differences in the behavioral output of the circuit. We measured the behavioral responses to vibration of early and late stage larvae in which the Chordotonal neurons were reversibly silenced during development (**Figure 7H**), just as in the functional imaging experiments described above (**Figure 7F**).

Early stage larvae in which Chordotonal neurons were reversibly silenced during embryonic development and later reactivated had a reduced probability to respond to vibration, compared to animals with unmanipulated Chordotonals (**Figure 7I-I’**). This behavioral defect persists in late stage larvae that had the same Chordotonal silencing period but grew for longer after activity was restored (**Figure 7J-J’**). These impaired behavioral responses could potentially be explained by the reduction in connectivity from silenced Chordotonal neurons onto Griddles and Ladders (**Figure 7D-E**), or onto some other neurons (not fully contained in the present EM volume). Particularly, reduced hunching responses (**Figure 7I’, J’**) are consistent with the reduced connectivity between Chordotonals and Griddles (**Figure 7D**), since Griddles have been shown to be required for hunching behavior (Jovanic et al., 2016). This indicates the connectivity alteration caused by the silencing of Chordotonal neurons has effects on the overall behavioral output of the circuit and persist even days after activity was restored.

### Basin cells compensate for the lack of mechanosensory input by increasing nociceptive input, a separate sensory modality

We found that upon silencing the mechanosensory Chordotonal neurons, Basin cells compensated by increasing their input from this silent sensory modality (**Figure 7B, G, G’**). We then asked whether Basins also compensate for the lack of mechanosensory input by increasing input from a different sensory modality. We have previously shown Basins are multisensory interneurons that receive input from both mechanosensory and nociceptive sensory neurons (Ohyama et al., 2015). We therefore looked at the connectivity between Basins and the nociceptive MD IV neurons revealed by EM reconstructions. We found that the fraction of Basin synapses from nociceptive neurons was greater when the mechanosensory Chordotonal neurons were silenced during development compared to when they were active (**Figure 8A**).

**Figure 8.**
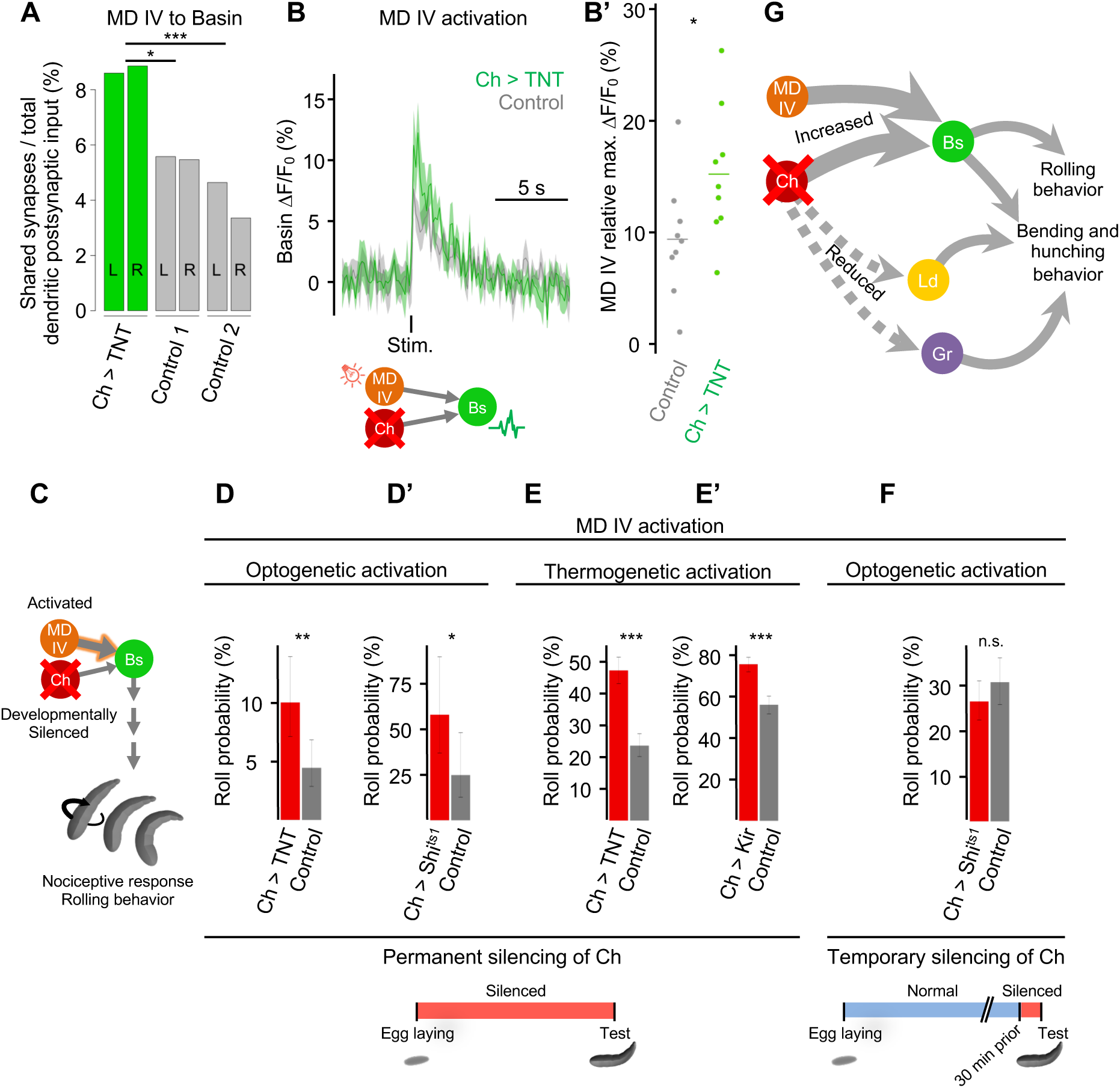
Basin cells compensate for the lack of mechanosensory input by increasing their nociceptive input. **A)** Connectivity from nociceptive MD IV onto Basin neurons revealed by EM reconstruction. The fraction of Basin dendritic input that is received from nociceptive MD IV increases when the Chordotonal (Ch) neurons are silenced by the targeted expression of TNT (green bars). Controls are two independent *w1118* animals (gray bars). The connectivity from the left (L) and right (R) sides of the nervous system is included to show consistency within sample. Left and right sides were grouped for statistical analysis using Chi-square test. **B)** Basin calcium responses (mean ± s.e.m.) to the optogenetic activation (Stim., 625 nm for 1 s) of nociceptive MD IV neurons increase when the Chordotonal cells are silenced by the targeted expression of TNT in third instar larvae (green trace; *Basin-GAL4 > UAS-GCaMP6s, MDIV-QF2 > QUAS-CsChrimson, Ch-LexA > LexAop-TNT*). Control responses (gray trace) are from animals lacking the TNT transgene (*Basin-GAL4 > UAS-GCaMP6s, MDIV-QF2 > QUAS-CsChrimson, Ch-LexA > +*). **B’)** Quantification of the calcium responses in B. Relative maximum ΔF/F0 values had the sample-specific baseline subtracted. n= 9 for each condition. Responses compared using Wilcoxon rank-sum test. **C)** Schematic of the experiment in which Chordotonal neurons were temporarily or permanently silenced, and the MD IV neurons were activated. Since the activation of MD IV normally elicits rolling behavior, this response was used to test for the behavioral effect of the increased connectivity between MD IV and Basins (A) generated by the inactivation of Chordotonals. **D-F)** Rolling behavior probabilities of third instar larvae to the activation of nociceptive MD IV neurons and permanent (D-E’) or temporary (F) silencing of Chordotonals. Error bars represent 95% confidence interval. **D-E’)** Optogenetic (D-D’) or thermogenetic (E-E’) activation of MD IV while Chordotonals were permanently silenced through various methods (using TNT for D and E; Shi^ts1^ for D’; Kir for E’) generated a higher probability of rolling behavior. **D)** Silencing of the Chordotonal neurons with the targeted expression of TNT (red bar; n= 298 animals) produced a higher rolling probability to the optogenetic activation of MD IV compared to control (gray bar; n= 426 animals). Genotypes were the same as in B. **D’)** Inactivation of Chordotonal neurons expressing Shi^ts1^ and optogenetic activation of MD IV led to increased rolling probability. Animals were incubated at 31 °C for three days, from egg laying until testing. Experimental animals (red) were *Ch-GAL4 > UAS-Shi^ts1^, MDIV-LexA > LexAop-CsChrimson*. Control animals (gray) were *+ > UAS-Shi^ts1^, MDIV-LexA > LexAop-CsChrimson*. Experimental n= 310 animals; control n= 322 animals. **E)** Silencing of the Chordotonal neurons with the targeted expression of TNT (red bar; n= 550 animals) produced a higher rolling probability to the thermogenetic activation of MD IV compared to control (gray bar; n= 526 animals). Experimental animals were *MDIV-LexA > LexAop-TrpA1, Ch-GAL4 > UAS-TNT*. Control animals were *MDIV-LexA > LexAop-TrpA1, + > UAS-TNT*. **E’)** Silencing of the Chordotonal neurons by the targeted expression of Kir (red bar; n= 580 animals) also led to an increased behavioral response, compared to control (gray bar; n= 512 animals). Experimental animals were *MDIV-LexA > LexAop-TrpA1, Ch-GAL4 > UAS-Kir*. Control animals were *MD IV-LexA > LexAop-TrpA1, + > UAS-Kir*. **F)** Temporary silencing of Chordotonal neurons shortly before (≥ 30 minutes before) and during the experiment with Shi^ts1^ has no effect on the rolling probability to the optogenetic activation of MD IV (red bar; n= 399 animals) compared to controls (gray bar; n= 305 animals). This suggests the connectivity compensation observed in A-B’ is due to a developmental effect in the circuit and not to the momentary effect of the loss of Chordotonal activity during the experiment. Experimental animals were *Ch-GAL4 > UAS-Shi^ts^, MD IV-LexA > LexAop-CsChrimson*. Control animals were *+ > UAS-Shi^ts^, MD IV-LexA > LexAop-CsChrimson*. **G)** Summary diagram of the connectivity effects of the developmental silencing of Chordotonal neurons. Basin cells (Bs) compensate for the lack of Chordotonal (Ch) input by increasing their input (thick arrow) from inactive (crossed out) Chordotonals. Additionally, Basins also show increased input from a separate sensory modality, the nociceptive MD IV neurons. This increase in connectivity has an effect on rolling behavior (D-F). However, the inhibitory Ladder (Ld) and Griddle (Gr) neurons lose (dashed arrow) Chordotonal input when Chordotonals were inactive.

In order to test whether these additional structural excitatory connections between nociceptive MD IV and Basins are functional, we measured Basin calcium responses to the activation of nociceptive neurons in animals in which the Chordotonal neurons are permanently silent. We simultaneously used CsChrimson to optogenetically activate MD IV neurons, GCaMP6s to monitor Basin activity, and TNT to constitutively silence Chordotonal neurons. Consistent with the EM connectivity data (**Figure 8A**), Basin cells have significantly greater calcium responses to nociceptive MD IV activation when the Chordotonal neurons were silenced than when they were intact (**Figure 8B-B’**).

EM reconstruction of structural connectivity and functional connectivity assays indicate Basins receive greater input from MD IV upon the absence of Chordotonal input (**Figure 8A-B’**). We therefore asked whether these increased connections have an effect on the behavioral output of the nociceptive circuit. Nociceptive MD IV and Basin neurons are part of the circuit underlying the rolling escape response of the larva, and the behavioral responses to their activation have been well characterized (Ohyama et al., 2015). Therefore, we monitored rolling escape behavioral responses to the activation of MD IV neurons in animals in which the Chordotonal neurons were silenced (**Figure 8C**).

We silenced Chordotonal neurons with TNT and activated MD IV neurons with CsChrimson (same genotype as in **Figure 8B**) and quantified their rolling behavioral responses. The rolling probability of these animals was significantly greater than those of animals in which the Chordotonals were not silenced (**Figure 8D**). As an alternative but equivalent approach, we silenced the Chordotonal neurons with Shi^ts1^ by rearing the animals at the restrictive temperature throughout their whole lives. These animals also displayed a significantly greater rolling probability (**Figure 8D’**). To further prove this effect, we used an additional alternative activation approach by thermogenetically activating the MD IV neurons. We selectively targeted the expression of TrpA1, a heat-activated cation channel (Hamada et al., 2008; Kang et al., 2012), in the nociceptive MD IV neurons. We thermogenetically activated MD IV neurons in animals in which the Chordotonals had been constitutively silenced by the targeted expression of TNT. These animals also displayed significantly greater rolling behavior probabilities (**Figure 8E**). In a separate experiment, we silenced the Chordotonal neurons with yet a third effector. We used Kir2.1 (Kir), an inwardly-rectifying potassium channel that hyperpolarizes the neurons and sets the resting membrane potential below the threshold required to fire action potentials (Baines et al., 2001; Johns et al., 1999). Silencing Chordotonals with Kir and thermogenetically activating MD IV with TrpA1 also generated increased rolling responses (**Figure 8E’**). We separately used four analogous manipulations to silence the Chordotonal neurons (with TNT, Shi^ts1^ or Kir) and used two different approaches to activate nociceptive MD IV neurons (optogenetic or thermogenetic). In each of these experiments, we observed an increase in nociceptive sensitivity as an effect of the lack of mechanosensory input (**Figure 8D-E’**). These results are consistent with the observed increase of structural and functional connections from nociceptive MD IV neurons onto Basins when Chordotonal neurons were silenced (**Figure 8A-B’**).

However, it is unclear whether this increase of nociceptive sensitivity is due to long-term developmental defects in the assembly of the Chordotonal circuit, or simply due to the immediate short-term effect of silencing Chordotonals during the experiment. Therefore, we silenced Chordotonals with Shi^ts1^ shortly before (≥30 minutes) and during the experiment (as opposed to silencing them constitutively) and measured the rolling behavioral responses to MD IV optogenetic activation. In this experiment, the behavioral responses are expected to be normal, since Chordotonals were only silenced briefly and the circuit’s connectivity could not have changed significantly. Indeed, silencing Chordotonals only during and briefly before testing generated no behavioral differences to MD IV activation compared to animals with intact Chordotonals (**Figure 8F**). Thus, the compensation in connectivity between nociceptive MD IV and Basin neurons, calcium responses, and behavior are all developmental effects of the silencing of Chordotonal neurons.

These results show that developmental activity may not play an instructive role in partner selectivity, but affects connectivity tuning with significant effects on circuit structure and behavior. Therefore, developmental activity is required for normal circuit formation.

## Discussion

Here we investigated whether partner-derived cues, global positional cues, and/or activity regulate 1) the selection of appropriate partners and 2) the appropriate numbers of connections formed with each partner during embryonic development using the tractable somatosensory circuitry of *Drosophila*. Specifically, we used EM reconstruction to determine whether selectively shifting or silencing one partner (mechanosensory Chordotonal neurons) prevents appropriate partner-recognition and/or the formation of appropriate numbers of connections with distinct types of postsynaptic partners. We then used functional connectivity and behavioral assays to relate the observed alterations in structural connectivity to functional connectivity and behavior.

During development, neurons follow global molecular guidance cues that instruct them to terminate in a specific 3D location of the nervous system (Couton et al., 2015;; Fukuhara et al., 2013;; Mauss et al., 2009;; Sürmeli et al., 2011;; Zlatic et al., 2003, 2009). It has been proposed that precise positioning of pre- and postsynaptic partners in the same location could reduce neuronal availability enough to generate specific connections (Balaskas et al., 2019; Li et al., 2007; Peters and Feldman, 1976; Rees et al., 2017; Roberts et al., 2014). Within this framework connectivity could result simply from promiscuous interactions in a delimited location where partner neurons independently converge, with no need for specific partner-recognition molecules. Alternatively, Sperry’s chemoaffinity hypothesis proposes that selective connections arise from specific partner-recognition molecules (Sperry, 1943, 1963). Evidence for both types of mechanisms have been found in different systems. On the one hand, a number of studies have shown that position of pre-and postsynaptic terminals can be specified independently of their partners by global positional cues (Couton et al., 2015; Mauss et al., 2009; Sürmeli et al., 2011; Zlatic et al., 2003, 2009). On the other hand, molecules have been identified in some systems that mediate partner matching, for example in the *Drosophila* visual system (Hong et al., 2012; Ward et al., 2015). Furthermore, previous light-microscopy studies in the olfactory system of *Drosophila* have shown that experimentally displaced sensory neurons are followed by their partner interneurons, preserving connectivity specificity (Li et al., 2019; Zhu et al., 2006). However, in these prior studies it was unclear how numbers of connections between partners and functional connectivity and behavior were affected when positional cues, or partner-recognition cues were altered.

Our results indicate that while developing sensory axons do use global non-partner derived positional cues to select their final termination area, partner-recognition molecules also exist in *Drosophila* nerve cord, and position alone does not specify connectivity. Thus, changing the precise location of mechanosensory neurons caused their postsynaptic partners to extend ectopic branches to reach for the displaced mechanosensory neurons and connect with them (**Figure 3** and **Figure 5**), preserving partner-partner specificity of the circuit. No new cell types were found to be strongly connected to shifted mechanosensory axons at their new location, which provides further evidence of remarkable partner specificity. Consistently, we found that animals in which mechanosensory axons were shifted to an ectopic location were still able to sense and respond to mechanosensory stimuli (**Figure 6I-J**) (but see below).

If partner-recognition molecules are sufficient for selective synaptogenesis irrespective of the location of partners, why is the precise location of sensory terminals so tightly regulated by the global non-partner derived positional cues? Our quantitative analysis of the numbers of synaptic inputs made by shifted sensory neurons onto specific postsynaptic partners provides a clue to this question. Thus, despite the fact that the postsynaptic neurons could find and connect to their presynaptic partners in ectopic locations, they did not manage to establish appropriate numbers of synaptic connections with them. Interestingly, when mechanosensory terminals were misplaced, some partners received less mechanosensory inputs than in controls, while others received more. Although these differences in connectivity may appear subtle, we observed significant differences in responses to mechanosensory stimuli in animals with misplaced mechanosensory terminals.

We do not know what causes this surprising quantitative difference in mechanosensory connections with different partners when mechanosensory neurons are shifted. One speculation is that not all neurons have equal capacity to search and reach for their partners, therefore not all manage to find them to the same extend. Another possibility is that the shift may cause a delay in partner-recognition and a delay in the formation of functional mechanosensory synaptic connections. This could in turn trigger network-level homeostatic plasticity mechanisms that alter the balance of excitation and inhibition within the circuit, resulting in increased numbers of connections onto excitatory interneurons, and reduced numbers of connections onto inhibitory interneurons. This latter possibility could explain why we observed similar quantitative effects on connectivity when mechanosensory neurons were shifted, and when they were inactivated during development (see below).

In many systems (especially in vertebrates), activity has been shown to play a major role in refining the patterns of connections between neurons during development (Kutsarova et al., 2017; Leighton and Lohmann, 2016; Tien and Kerschensteiner, 2018). Landmark studies in kittens have shown that depriving them of visual input during an early critical period by suturing their eyelids permanently impaired their vision, even after their eyes were opened again (Hubel and Wiesel, 1964). Activity can refine the patterns of connections through Hebbian and/or homeostatic plasticity mechanisms (Kaneko et al., 2017; Marder, 2011; Schulz and Lane, 2017; Sheng et al., 2018; Sugie et al., 2018; Tien and Kerschensteiner, 2018; Tripodi et al., 2008; Turrigiano, 2017; Yuan et al., 2011). However, what kinds of changes are induced within the network in response to selective silencing of specific neuron types is not fully understood. Specifically, the extent to which silencing of a specific neuron modulates the structural connections and synapse numbers within the circuit as opposed to only functional connections is still unclear.

In the insect central nervous system, the role of activity in refining synaptic connectivity is less well established. Several studies have shown that lack of sensory activity during development does not affect neuron morphology or the capacity to form connections (Baines et al., 2001; Constance et al., 2018; Hiesinger et al., 2006; Jefferis et al., 2004; Scott et al., 2003). At the same time, other studies have reported neural circuits can adapt their morphology, connectivity, and behavior in response to changes in developmental activity (Giachello and Baines, 2015, 2017; Kaneko et al., 2017; Prieto-Godino et al., 2012; Sheng et al., 2018; Tripodi et al., 2008; Wolfram and Baines, 2013; Yuan et al., 2011). Furthermore, most of these studies lack a comprehensive synaptic-resolution analysis of the effects of silencing a specific neuron type on the numbers of connections between all partners. Thus, the role that activity plays in *Drosophila* is still an open question.

Our EM reconstructions revealed that silenced mechanosensory neurons connected to appropriate partners, but they formed inappropriate numbers of connections. Interestingly, excitatory interneurons (Basin) received a higher fraction of input from silenced mechanosensory neurons compared to controls, while the inhibitory interneurons (Ladder, Griddle, and Drunken) received a lower fraction of mechanosensory input. We also found that selective silencing of mechanosensory neurons led to an increase in the fraction of nociceptive neuron input onto multimodal excitatory Basins (**Figure 8**). This overall effect is reminiscent to observations in the cortex where sensory deprivation induces network-level homeostasis that alters the balance of excitation and inhibition within the network (Maffei et al., 2004, Mendelsohn et al., 2015). Furthermore, synaptic scaling in the cortex is thought to be multiplicative, such that all excitatory connections onto an excitatory neuron are scaled by the same amount when excitatory drive onto that neuron is reduced (Turrigiano and Nelson, 2004). In contrast, the inhibitory connections onto excitatory neurons are reduced. While the majority of studies in cortex focus on homeostatic plasticity of functional connections, we observe a drastic plasticity in the number of synaptic connections between partners. This apparent homeostasis of synapse numbers may also follow similar multiplicative rules, because we found that both mechanosensory and nociceptive inputs onto excitatory multimodal interneurons (Basin) were increased when only mechanosensory neurons were selectively silenced.

We also observed changes in functional connectivity within the network that were consistent with the observed changes in structural connectivity. Furthermore, we found that selective silencing of mechanosensory neurons, resulted in alterations in stimulus-specific behavioral responses. Larvae with permanently silenced mechanosensory neurons showed increased behavioral responses to nociceptive stimuli, consistent with the observed increased number of connections from nociceptive neurons onto downstream multimodal Basin interneurons, as well as with the increase in functional connection strength between them. This structural and behavioral compensation is reminiscent to what has been observed in mammals, where if one sensory modality is removed, another modality is restructured and improved (Lomber et al., 2010; Rauschecker, 1995; Rauschecker and Korte, 1993).

Interestingly, we also found that larvae in which mechanosensory neurons were selectively silenced only during development had permanently decreased responses to mechanosensory stimuli, even days after activity was restored. This behavioral result is also reminiscent to the findings in mammals, where deprivation of visual input during an early critical period permanently impairs vision (Hubel and Wiesel, 1964), however it appears at odds with the increased number of mechanosensory connections and increased functional connection strength from mechanosensory neurons onto the postsynaptic multimodal Basin neurons. A possible explanation for this is the observed reduction in the fraction of mechanosensory connections onto inhibitory neurons. Unlike the nociceptive neurons, the mechanosensory neurons have more inhibitory than excitatory postsynaptic partners, and these inhibitory interneurons contribute to mechanosensory behaviors through disinhibition (Jovanic et al., 2016). Silencing the mechanosensory neurons may therefore result in a permanent reduction in disinhibition with permanent consequences on behavior.

In summary, while partner-recognition molecules can ensure that neurons recognize and connect only with appropriate partners, they are not sufficient to robustly specify appropriate numbers of synapses with distinct postsynaptic partners. Conversely, while neither precise location of presynaptic terminals, nor neuronal activity in presynaptic partners directly instructs partner specificity, both are crucial to achieve appropriate numbers of connections with distinct postsynaptic partners, appropriate strength of functional connections, and appropriate behavior. Furthermore, we find that position and activity influence connections onto excitatory and inhibitory interneurons in opposite ways. To our knowledge, our study reveals with unprecedented detail the fine connectivity effects of neuron location, identity, and activity on synaptic specificity, showcasing the role of multiple factors that must work together to influence circuit formation, in a highly cell-type specific way.

## Materials and methods

### Fly stocks

All animals used in this study are of the *Drosophila melanogaster* species and were kept on fly food at 25 °C unless otherwise specified. The fly food composition is as follows: molasses 5.1% v/v, dry yeast 2.04% m/v, corn meal 8.45% m/v, agar 0.75% m/v, Tegosept 0.2% v/v, and propionic acid 0.5 % v/v. Animals for optogenetic experiments were kept in the dark on fly food supplemented with all-trans-retinal (Cat. #R240000, Toronto Research Chemicals) to a concentration of 0.5 mM.

All throughout this document, abbreviated names of the fly strains have been used for simplicity. Different driver lines were used to restrict the expression of a given transgene to the neurons of interest. The GAL4/UAS, LexA/LexAop, and QF/QUAS binary expression systems (del Valle Rodríguez et al., 2012) were used interchangeably. The specific expression system used for each experiment is stated where appropriate.

The *R72F11* driver was used for transgene expression in Basin cells (Ohyama et al., 2015), *iav* or *R61D08* for Chordotonal mechanosensory neurons (Kwon et al., 2010; Ohyama et al., 2015), *ppk* for multidendritic class IV neurons (Ainsley et al., 2003), and *R71A10* for A00c neurons (Ohyama et al., 2015). The *w;; attP2* line has an empty insertion site with no driver and was used as control for some experiments (where indicated).

### Live imaging

For live imaging experiments, fly stocks were generated to label Basin cells with myristoylated GFP using the *72F11-LexA* driver, and the Chordotonal sensory neurons with myristoylated tdTomato using the *iav-GAL4* driver. These animals contained a mutation in the myosin heavy chain (*mhc*[*1*]) that disables muscle contraction in homozygous mutants in order to prevent interruptions during the imaging process (Mogami and Hotta, 1981; O’Donnell and Bernstein, 1988; Vonhoff and Keshishian, 2017). This mutation was kept over the balancer CyO to establish viable stocks. When possible, CyO labelled with dfd-GMR Yellow fluorescent protein (DGY) (Le et al., 2006) was used to facilitate the selection of homozygous embryos. For the live imaging of Basins only, the following line was used: *w; R72F11-LexA, LexAop-GFP, mhc*[*1*]*/CyO, DGY; iav-GAL4, UAS-tdT*. For simultaneous live imaging of Basins and Chordotonals, the following line was used: *w; R72F11-LexAp65 in JK22C, 13XLexAop2-IVS-myr::GFP in su(Hw)attP5, mhc*[*1*]*/CyO, DGY; iav-GAL4, UAS-IVS-myr::tdTomato in attP2*.

Eggs were collected for one hour at 25 °C on agar plates with yeast paste. After collection, the eggs were incubated at 25 °C for 13 hours. Then the eggs were treated with a 1:1 mixture of water and commercial bleach for five minutes or until the chorion was fully removed. The resulting mixture was passed through a sieve to recover the dechorionated eggs. These were rinsed with distilled water to remove bleach and transferred into a Petri dish. Single embryos were carefully picked under a dissection microscope and placed ventral side up on an oxygen-permeable teflon membrane (Lumox). Such membrane was stretched on a custom-made mount that can hold liquid and fits the microscope stage. Multiple embryos were aligned in a row and fully covered with room temperature distilled water. This was done not more than 10 minutes after the embryos were dechorionated to prevent dehydration.

The imaging setup consisted of a Yokogawa CSU-22 spinning disk confocal field scanner mounted on an Olympus BX51 WI fixed-stage upright compound microscope, with an Evolve EMCCD camera (Photometrics) and a LUMPlanFl 60X/0.9 NA (Olympus) water dipping objective. The excitation wavelengths for imaging GFP and tdT were 488 nm and 561 nm, respectively. 50 μm Z-stacks with a 1 μm step size and 218 nm/pixel resolution were acquired in two imaging channels every time point for each embryo. Multiple embryos were imaged one after the other continuously for at least 12 hours. The time point frequency varied from 1 to 5 min depending on the number of embryos imaged simultaneously in each session. The center of the stack in the Z axis was roughly located at the center of the developing ventral nerve cord at the beginning of the imaging session. The imaging range in the Z axis was manually readjusted during the session if needed to ensure coverage of the neurons of interest. The images were acquired with the control of MetaMorph software (Molecular devices).

### Image processing for live imaging data

Standard image processing was performed using Fiji (Rueden et al., 2017; Schindelin et al., 2012). Briefly, the imaging stacks were cropped to remove Z sections that did not contain the neurons of interest. The images were denoised using nd-safir (Boulanger et al., 2010). Z-projections were generated, and the imaging channels were merged to create 2D time-lapse videos of the developing neurons in two colors. Bleach correction (Fiji) was used to adjust for the increasing brightness of the neurons through time. Ilastik (Sommer et al., 2011) was used for pixel classification to generate the segmented images. Different trained pixel classification parameters were used for each imaging channel.

### Calcium imaging with GCaMP

Calcium responses were imaged as GCaMP6s (Chen et al., 2013) fluorescence fluctuations in the neurons of interest (Basin or A00c). CsChrimson was expressed in presynaptic neurons (Chordotonal or MD IV) for optogenetic activation (Klapoetke et al., 2014). GCaMP signals were recorded in dissected central nervous systems in a saline solution (135 mM NaCl, 5 mM KCl, 2 mM CaCl2×·2H2O, 4 mM MgCl2×·6H2O, 5 mM TES, 36 mM Sucrose, pH 7.15) and adhered by the ventral side to a cover glass coated with poly-L-lysine (SIGMA, P1524) on a small Sylgard (Dow Corning) plate.

The calcium imaging experiments were performed using a 3i VIVO Multiphoton upright microscope (Intelligent Imaging Innovations). The Chordotonal neurons were photo-stimulated using a 1040 nm laser (1040-3 femtoTrain, Spectra-Physics) coupled to a 2-photon Phasor (Intelligent Imaging Innovations) to generate a holographic pattern to restrict the activation area. GCaMP responses were recorded using an imaging laser tuned to 925 nm (Insight DS+ Dual, Spectra-Physics) and an Apo LWD 25x/1.10W objective (Nikon).

For the reversible silencing of Chordotonal neurons with Shibire^ts1^ (Chen et al., 1991) and recording of Basin responses the *w; R61D08-LexA; R72F11-GAL4* line was crossed to: *w; LexAop-Shi; UAS-GCaMP6s, LexAop-CsChrimson* for experimental animals, or to *w;; UAS-GCaMP6s, LexAop-CsChrimson* for control. Embryos were collected on retinal food for two hours at 25 °C and then incubated in the dark at 31 °C for 24 hours, and for another day at 18 °C until testing. For the activation of Chordotonal neurons and recording of Basin responses, the stimulation protocol consisted of an initial 30 s resting period, a 100 ms stimulation event, and a final 30 s resting period. A photo-stimulation region of 26.3 μm × 11.9 μm was delimited to contain the Chordotonal axon terminals within one abdominal hemisegment, approximately. The stimulation power value measured at the objective end with a power meter (PM100D Thorlabs) was 34.2 mW. This protocol was executed in three different abdominal hemisegments per sample. Any two stimulated ipsilateral hemisegments were separated by at least one unstimulated hemisegment as a precaution in case of unintended leaky stimulation of the adjacent hemisegment. GCaMP responses were imaged at the Basin axons on a single Z plane at 6.61 frames/s.

For the activation of Chordotonal neurons and imaging of A00c calcium responses, the *w; R61D08-LexA; R71A10-GAL4* line was crossed to: *w; LexAop-Shi; UAS-GCaMP6s, LexAop-CsChrimson* for experimental animals, or to *w;; UAS-GCaMP6s, LexAop-CsChrimson* for control. A photo-stimulation region of 16.2 μm × 56.9 μm was set to cover the Chordotonal axons in the most anterior thoracic hemisegments. The protocol consisted an initial resting period of 30 s, a stimulation event of 200 ms and 323 mW, and a final resting period of 30 s. This protocol was implemented twice per sample, stimulating Chordotonal axons on either side of the nerve cord, one at a time. A00c calcium responses were recorded at their axons in the corresponding side of the brain at 8.79 frames/s.

For the experiments where the Chordotonal neurons were silenced with tetanus toxin (TNT) (Sweeney et al., 1995) and the MD IV neurons were optogenetically activated, the *w; R61D08-LexA; R72F11-GAL4, ppk-QF2* line was crossed to: *w, QUAS-CsChrimson; LexAop-TNT; UAS-GCaMP6s* for experimental animals, or to *w, QUAS-CsChrimson;; UAS-GCaMP6s* for control animals. Eggs were collected on retinal food and incubated in the dark at 25°C for four days. The dissected samples were left in the dark for at least two minutes immediately before initiating the imaging session. All the MD IV axons were photo-stimulated with a 625 nm LED mounted on the microscope stage to illuminate the entire sample with 170 μW/cm^2^. The stimulation protocol consisted of an initial 30 s resting period, four 1 s stimulation events of the same intensity, each followed by a 30 s resting period. This protocol was executed once per sample. All other imaging details are as stated above.

### Image analysis of calcium imaging data

The GCaMP image data were processed using custom macros in Fiji (Schindelin et al., 2012) and analyzed using custom code written in R (R Core Team, 2015). Briefly, a region of interest (ROI) was manually defined to include the corresponding GCaMP-expressing axons. The average pixel value inside such ROI was measured with Fiji across all time points for each sample. All fluorescence values were reported relative to a fluorescence baseline (F0) defined as the median pixel value of the corresponding ROI during the entire imaging experiment. ΔF/F0 was calculated as ΔF/F0= (Ft – F0)/F0, where Ft is the mean fluorescence value of the ROI at a given time point. The relative maximum ΔF/F0 was defined as the maximum ΔF/F0 value in a 4.5 s time window immediately after stimulation offset from which the baseline (mean ΔF/F0 of the 3 s preceding stimulation onset) was subtracted. Those individual trials in which there were no responses were discarded. A trial with no response was defined as that where the mean ΔF/F0 in the 4.5 s following stimulation was within ±1.5 (for Chordotonal activation) or ±0.5 (for MD IV activation) standard deviations of the baseline (3 s preceding stimulation). Individual imaging trials were averaged by animal. The calcium imaging data were plotted using the ggplot2 (Wickham, 2009) package in R.

### Behavioral assays

All the behavioral apparatuses used in this study have been described previously (Ohyama et al., 2013, 2015) and will only be explained briefly. The rigs had some common core components and differed mostly in the hardware to deliver different types of stimuli. Generally, all consisted of a temperature-controlled enclosure with a high-resolution camera on top, an array of infrared (850 nm) LEDs for illumination, a computer for data acquisition and storage, and the respective hardware modules to deliver and control different stimuli.

For thermogenetic activation, the neurons of interest expressed the heat-activated cation channel TrpA1 (Hamada et al., 2008; Kang et al., 2012). For these experiments, eggs were collected on food plates for 6-8 hours and incubated at 18 °C for 8 days, unless otherwise stated. The animals were placed on a thin layer of 4% charcoal agar on top of an aluminum plate. This was placed on a Peltier module to control temperature to the desired value. The standard thermogenetic activation protocol consisted of 30 s at 20 °C, followed by a ramping-up period of 40 s to reach 35 °C, 50 s at 35 °C, and a final ramping-down period of 60 s to reach 20 °C. Whenever optogenetic activation was paired with a thermal stimulus, red (630 nm) LEDs were used with a power density of 490 μW/cm^2^ onto the center of the plate.

For vibration experiments, eggs were collected on food plates for 6-8 hours and incubated at 25°C for four days, unless otherwise stated. The mechanical stimulus was delivered as vibration using a speaker located to the side of a 4% agar plate holding the animals. Tones were played at 1000 Hz, with a measured volume (Extech, 407730) of 122 dB. The protocol consisted of 30 s of no sound, 30 s tone at 1000 Hz, and 30 s of no sound.

For optogenetic activation, animals carried the CsChrimson transgene (Klapoetke et al., 2014) in the neurons of interest. Eggs were collected on retinal food for 6-8 hours and incubated in the dark at 25 °C for four days, unless otherwise specified. When photo-activation was the only stimulus, larvae were placed on a 4% agar plate on top of an array of red (630 nm) LEDs with power density of 638 μW/cm^2^ through the plate. The activation protocol consisted of 30 s of the LEDs being off, 15 s on, 30 s off, 15 s on, and 30 s off.

For each behavioral experiment, a total of roughly 400-500 animals were tested across multiple trials. For experiments performed on a thermal plate, each trial included approximately 20 animals. All other experimental trials included approximately 50 animals each. The number of animals from experiments that included young (before 3^rd^ instar) larvae is much lower due to technical difficulties of handling and tracking smaller animals. Many animal traces are discarded throughout the subsequent analysis pipeline. The resulting number of animals used for statistical analysis varies across experiments and depends on the nature of the behavior evoked, stimulus and size of behavioral plate.

Stimulus control, object detection, and feature extraction were performed by the Multi Worm Tracker and SALAM-LARA (http://sourceforge.net/projects/salam-hhmi) software as previously described (Denisov et al., 2013; Ohyama et al., 2013).

### Electron microscopy reconstruction

Four electron microscopy volumes were used in this study. They comprise a whole or partial central nervous system of first instar *Drosophila* larvae. Two of these are control volumes of a *w1118* genotype and have been previously reported (Ohyama et al., 2015). The neurons from the two control volumes were previously reconstructed by members and collaborators of the Cardona lab (Janelia Research Campus, HHMI). The two remaining EM volumes were acquired for this study using the same protocol reported for the control volumes (Ohyama et al., 2015). They have an image resolution of 3.8 nm by 3.8 nm by 40 nm in x, y and z, respectively. These volumes include a 1.5-segment fraction of the central nervous system (A2 and A3 segments) of first instar larvae. The genotypes for these volumes are: 1) *w;; iav-GAL4/UAS-FraRobo* 2) *w; UAS-TNT/+; iav-GAL4/+*. The neurons were reconstructed using CATMAID (Saalfeld et al., 2009) to obtain the skeletonized structure and connectivity of the cells of interest. All the connectivity data were generated in CATMAID and processed in R.

### Statistical analysis

Statistical analysis was performed using R. The calcium responses between control and experimental animals were compared using the single-sided Wilcoxon rank-sum test.

For the behavioral assays, the probability of a behavior occurring was calculated as the proportion of animals that performed the specified behavior at least once during the 15 s (for optogenetic activation or vibration stimulus) or 40 s (for thermogenetic activation) immediately after stimulus onset across all trials. The analysis time window for thermogenetic activation is longer due to its slower activation resulting from temperature ramping. Therefore, the stimulus onset for thermogenetic activation experiments was defined as the moment the thermal plate reached 35 °C. Only those animals that were detected for at least 95% of the analyzed time window and did not contacted another animal during this period were included in the analysis. The behavior probabilities were compared using a chi-square test for proportions. Behavior durations were calculated for the time windows mentioned above and compared using a double-sided t-test.

Electron microscopy connectivity data were compared using a chi-square test for proportions.

In all figures, * represents p-value ≤ 0.05, ** represents p-value ≤ 0.01, and *** represents p-value ≤ 0.001.

### Immunohistochemistry

Larval brains were dissected in PBS, mounted on 12mm #1.5 thickness poly-L-lysine coated coverslips (Neuvitro Corporation, Vancouver, WA, Cat# H-12-1.5-PLL) and fixed for 23 minutes in fresh 4% paraformaldehyde (PFA) (Electron Microscopy Sciences, Hatfield, PA, Cat. 15710) in PBST. Brains were washed in PBST and then blocked with 2.5% normal donkey serum and 2.5% normal goat serum (Jackson ImmunoResearch Laboratories, Inc., West Grove, PA) in PBST overnight. Brains were incubated in primary antibody for two days at 4°C. The primary was removed and the brains were washed with PBST, then incubated in secondary antibodies overnight at 4°C. The secondary antibody was removed following overnight incubation and the brains were washed in PBST. Brains were dehydrated with an ethanol series (30%, 50%, 75%, 100%, 100%, 100% ethanol; all v/v, 10 minutes each) (Decon Labs, Inc., King of Prussia, PA, Cat. 2716GEA) then incubated in xylene (Fisher Chemical, Eugene, OR, Cat. X5-1) for 2x 10 minutes. Samples were mounted onto slides containing DPX mountant (Millipore Sigma, Burlington, MA, Cat. 06552) and cured for 3 days then stored at 4°C until imaged.

## References

Ainsley, J.A., Pettus, J.M., Bosenko, D., Gerstein, C.E., Zinkevich, N., Anderson, M.G., Adams, C.M., Welsh, M.J., and Johnson, W.A. (2003). Enhanced Locomotion Caused by Loss of the Drosophila DEG/ENaC Protein Pickpocket1. Curr. Biol. 13, 1557–1563.

Akin, O., Bajar, B.T., Keles, M.F., Frye, M.A., Lawrence, S., Correspondence, Z., and Lawrence Zipursky, S. (2019). Cell-type-Specific Patterned Stimulus-Independent Neuronal Activity in the Drosophila Visual System during Synapse Formation. Neuron 101, 894–904.

Araújo, S.J., and Tear, G. (2003). Axon guidance mechanisms and molecules: lessons from invertebrates. Nat. Rev. Neurosci. 4, 910–922.

Baines, R.A., and Bate, M. (1998). Electrophysiological development of central neurons in the Drosophila embryo. J. Neurosci. 18, 4673–4683.

Baines, R.A., Uhler, J.P., Thompson, A., Sweeney, S.T., and Bate, M. (2001). Altered electrical properties in Drosophila neurons developing without synaptic transmission. J. Neurosci. 21, 1523–1531.

Balaskas, N., Abbott, L.F., Jessell, T.M., and Ng, D. (2019). Positional Strategies for Connection Specificity and Synaptic Organization in Spinal Sensory-Motor Circuits. Neuron 0.

Bashaw, G.J., and Goodman, C.S. (1999). Chimeric axon guidance receptors: the cytoplasmic domains of slit and netrin receptors specify attraction versus repulsion. Cell 97, 917–926.

Boulanger, J., Kervrann, C., Bouthemy, P., Elbau, P., Sibarita, J.-B., and Salamero, J. (2010). Patch-Based Nonlocal Functional for Denoising Fluorescence Microscopy Image Sequences. IEEE Trans. Med. Imaging 29, 442–454.

Brown, T.H., Zhao, Y., and Leung, V. (2009). Hebbian Plasticity. Encycl. Neurosci. 1049–1056.

Chen, M.S., Obar, R.A., Schroeder, C.C., Austin, T.W., Poodry, C.A., Wadsworth, S.C., and Vallee, R.B. (1991). Multiple forms of dynamin are encoded by shibire, a Drosophila gene involved in endocytosis. Nature 351, 583–586.

Chen, T.-W., Wardill, T.J., Sun, Y., Pulver, S.R., Renninger, S.L., Baohan, A., Schreiter, E.R., Kerr, R.A., Orger, M.B., Jayaraman, V., et al. (2013). Ultrasensitive fluorescent proteins for imaging neuronal activity. Nature 499, 295–300.

Constance, W.D., Mukherjee, A., Fisher, Y.E., Pop, S., Blanc, E., Toyama, Y., and Williams, D.W. (2018). Neurexin and Neuroligin-based adhesion complexes drive axonal arborisation growth independent of synaptic activity. Elife 7.

Couton, L., Mauss, A.S., Yunusov, T., Diegelmann, S., Evers, J.F., and Landgraf, M. (2015). Development of Connectivity in a Motoneuronal Network in Drosophila Larvae. Curr. Biol. 25, 568–576.

Denisov, G., Ohyama, T., Jovanic, T., and Zlatic, M. (2013). Model-based Detection and Analysis of Animal Behaviors using Signals Extracted by Automated Tracking. In BIOSIGNALS, p.

Dickson, B.J. (2002). Molecular mechanisms of axon guidance. Science 298, 1959–1964.

Eichler, K., Li, F., Litwin-Kumar, A., Park, Y., Andrade, I., Schneider-Mizell, C.M., Saumweber, T., Huser, A., Eschbach, C., Gerber, B., et al. (2017). The complete connectome of a learning and memory centre in an insect brain. Nature 548, 175–182.

Fukuhara, K., Imai, F., Ladle, D.R., Katayama, K., Leslie, J.R., Arber, S., Jessell, T.M., and Yoshida, Y. (2013). Specificity of Monosynaptic Sensory-Motor Connections Imposed by Repellent Sema3E-PlexinD1 Signaling. Cell Rep. 5, 748–758.

Gerhard, S., Andrade, I., Fetter, R.D., Cardona, A., and Schneider-Mizell, C.M. (2017). Conserved neural circuit structure across Drosophila larval development revealed by comparative connectomics. Elife 6, e29089.

Giachello, C.N., and Baines, R.A. (2015). Inappropriate Neural Activity during a Sensitive Period in Embryogenesis Results in Persistent Seizure-like Behavior. Curr. Biol. 25, 2964–2968.

Giachello, C.N., and Baines, R.A. (2017). Regulation of motoneuron excitability and the setting of homeostatic limits. Curr. Opin. Neurobiol. 43, 1–6.

Hamada, F.N., Rosenzweig, M., Kang, K., Pulver, S.R., Ghezzi, A., Jegla, T.J., and Garrity, P.A. (2008). An internal thermal sensor controlling temperature preference in Drosophila. Nature 454, 217–220.

Helmstaedter, M., Briggman, K.L., Turaga, S.C., Jain, V., Seung, H.S., and Denk, W. (2013). Connectomic reconstruction of the inner plexiform layer in the mouse retina. Nature 500, 168–174.

Hiesinger, P.R., Zhai, R.G., Zhou, Y., Koh, T.W., Mehta, S.Q., Schulze, K.L., Cao, Y., Verstreken, P., Clandinin, T.R., Fischbach, K.F., et al. (2006). Activity-Independent Prespecification of Synaptic Partners in the Visual Map of Drosophila. Curr. Biol. 16, 1835–1843.

Hong, W., and Luo, L. (2014). Genetic control of wiring specificity in the fly olfactory system. Genetics 196, 17–29.

Hong, W., Mosca, T.J., and Luo, L. (2012). Teneurins instruct synaptic partner matching in an olfactory map. Nature 484, 201–207.

Hubel, D.H., and Wiesel, T.N. (1964). EFFECTS OF MONOCULAR DEPRIVATION IN KITTENS. Naunyn. Schmiedebergs. Arch. Exp. Pathol. Pharmakol. 248, 492–497.

Jefferis, G.S.X.E., Vyas, R.M., Berdnik, D., Ramaekers, A., Stocker, R.F., Tanaka, N.K., Ito, K., and Luo, L. (2004). Developmental origin of wiring specificity in the olfactory system of Drosophila. Development 131, 117–130.

Jenett, A., Rubin, G.M., Ngo, T.-T.B., Shepherd, D., Murphy, C., Dionne, H., Pfeiffer, B.D., Cavallaro, A., Hall, D., Jeter, J., et al. (2012). A GAL4-Driver Line Resource for Drosophila Neurobiology. Cell Rep. 2, 991–1001.

Johns, D.C., Marx, R., Mains, R.E., O’Rourke, B., and Marbán, E. (1999). Inducible genetic suppression of neuronal excitability. J. Neurosci. 19, 1691–1697.

Jovanic, T., Schneider-Mizell, C.M., Shao, M., Masson, J.-B., Denisov, G., Fetter, R.D., Mensh, B.D., Truman, J.W., Cardona, A., and Zlatic, M. (2016). Competitive Disinhibition Mediates Behavioral Choice and Sequences in Drosophila. Cell 167, 858–870.e19.

Kaneko, T., and Ye, B. (2015). Fine-scale topography in sensory systems: insights from Drosophila and vertebrates. J. Comp. Physiol. A 201, 911–920.

Kaneko, T., Macara, A.M., Li, R., Hu, Y., Iwasaki, K., Dunnings, Z., Firestone, E., Horvatic, S., Guntur, A., Shafer, O.T., et al. (2017). Serotonergic Modulation Enables Pathway-Specific Plasticity in a Developing Sensory Circuit in Drosophila. Neuron 95, 623–638.e4.

Kang, K., Panzano, V.C., Chang, E.C., Ni, L., Dainis, A.M., Jenkins, A.M., Regna, K., Muskavitch, M.A.T., and Garrity, P.A. (2012). Modulation of TRPA1 thermal sensitivity enables sensory discrimination in Drosophila. Nature 481, 76–80.

Kasthuri, N., Hayworth, K.J., Berger, D.R., Schalek, R.L., Conchello, J.A., Knowles-Barley, S., Lee, D., Vázquez-Reina, A., Kaynig, V., Jones, T.R., et al. (2015). Saturated Reconstruction of a Volume of Neocortex. Cell 162, 648–661.

Kilman, V., van Rossum, M.C.W., and Turrigiano, G.G. (2002). Activity deprivation reduces miniature IPSC amplitude by decreasing the number of postsynaptic GABA(A) receptors clustered at neocortical synapses. J. Neurosci. 22, 1328–1337.

Kitamoto, T. (2001). Conditional modification of behavior inDrosophila by targeted expression of a temperature-sensitiveshibire allele in defined neurons. J. Neurobiol. 47, 81–92.

Klapoetke, N.C., Murata, Y., Kim, S.S., Pulver, S.R., Birdsey-Benson, A., Cho, Y.K., Morimoto, T.K., Chuong, A.S., Carpenter, E.J., Tian, Z., et al. (2014). Independent optical excitation of distinct neural populations. Nat. Methods 11, 338–346.

Kolodkin, A.L., and Tessier-Lavigne, M. (2011). Mechanisms and molecules of neuronal wiring: a primer. Cold Spring Harb. Perspect. Biol. 3.

Krishnaswamy, A., Yamagata, M., Duan, X., Hong, Y.K., and Sanes, J.R. (2015). Sidekick 2 directs formation of a retinal circuit that detects differential motion. Nature 524, 466–470.

Kutsarova, E., Munz, M., and Ruthazer, E.S. (2017). Rules for Shaping Neural Connections in the Developing Brain. Front. Neural Circuits 10, 111.

Kwon, Y., Shen, W.L., Shim, H.-S., and Montell, C. (2010). Fine thermotactic discrimination between the optimal and slightly cooler temperatures via a TRPV channel in chordotonal neurons. J. Neurosci. 30, 10465–10471.

Langley, J.N. (1895). Note on Regeneration of Prae-Ganglionic Fibres of the Sympathetic. J. Physiol. 18, 280–284.

Le, T., Liang, Z., Patel, H., Yu, M.H., Sivasubramaniam, G., Slovitt, M., Tanentzapf, G., Mohanty, N., Paul, S.M., Wu, V.M., et al. (2006). A New Family of Drosophila Balancer Chromosomes With a w-dfd-GMR Yellow Fluorescent Protein Marker. Genet. Soc. Am.

Lee, W.-C.A., Bonin, V., Reed, M., Graham, B.J., Hood, G., Glattfelder, K., and Reid, R.C. (2016). Anatomy and function of an excitatory network in the visual cortex. Nature 532, 370–374.

Leighton, A.H., and Lohmann, C. (2016). The Wiring of Developing Sensory Circuits—From Patterned Spontaneous Activity to Synaptic Plasticity Mechanisms. Front. Neural Circuits 10, 71.

Li, H., Li, T., Horns, F., Li, J., Xie, Q., Xu, C., Wu, B., Kebschull, J.M., Vacek, D., Xie, A., et al. (2019). Coordinating Receptor Expression and Wiring Specificity in Olfactory Receptor Neurons. BioRxiv 594895.

Li, W.-C., Cooke, T., Sautois, B., Soffe, S.R., Borisyuk, R., and Roberts, A. (2007). Axon and dendrite geography predict the specificity of synaptic connections in a functioning spinal cord network. Neural Dev. 2, 17.

Lomber, S.G., Meredith, M.A., and Kral, A. (2010). Cross-modal plasticity in specific auditory cortices underlies visual compensations in the deaf. Nat. Neurosci. 13, 1421–1427.

Maffei, A., and Turrigiano, G.G. (2008). Multiple modes of network homeostasis in visual cortical layer 2/3. J. Neurosci. 28, 4377–4384.

Maffei, A., Nelson, S.B., and Turrigiano, G.G. (2004). Selective reconfiguration of layer 4 visual cortical circuitry by visual deprivation. Nat. Neurosci. 7, 1353–1359.

Marder, E. (2011). Variability, compensation, and modulation in neurons and circuits. Proc. Natl. Acad. Sci. U. S. A. 108 Suppl 3, 15542–15548.

Mauss, A., Tripodi, M., Evers, J.F., and Landgraf, M. (2009). Midline Signalling Systems Direct the Formation of a Neural Map by Dendritic Targeting in the Drosophila Motor System. PLoS Biol 7, 1000200.

Mendelsohn, A.I., Simon, C.M., Abbott, L.F., Mentis, G.Z., and Jessell, T.M. (2015). Activity Regulates the Incidence of Heteronymous Sensory-Motor Connections. Neuron 87, 111–123.

Mogami, K., and Hotta, Y. (1981). Isolation of Drosophila flightless mutants which affect myofibrillar proteins of indirect flight muscle. Mol. Gen. Genet. 183, 409–417.

O’Donnell, P.T., and Bernstein, S.I. (1988). Molecular and ultrastructural defects in a Drosophila myosin heavy chain mutant: differential effects on muscle function produced by similar thick filament abnormalities. J. Cell Biol. 107, 2601–2612.

Ohyama, T., Jovanic, T., Denisov, G., Dang, T.C., Hoffmann, D., Kerr, R.A., and Zlatic, M. (2013). High-Throughput Analysis of Stimulus-Evoked Behaviors in Drosophila Larva Reveals Multiple Modality-Specific Escape Strategies. PLoS One 8, e71706.

Ohyama, T., Schneider-Mizell, C.M., Fetter, R.D., Valdes-Aleman, J., Franconville, R., Rivera-Alba, M., Mensh, B.D., Branson, K.M., Simpson, J.H., Truman, J.W., et al. (2015). A multilevel multimodal circuit enhances action selection in Drosophila. Nature 520, 633–639.

Peters, A., and Feldman, M.L. (1976). The projection of the lateral geniculate nucleus to area 17 of the rat cerebral cortex. I. General description. J. Neurocytol. 5, 63–84.

Pfeiffer, B.D., Jenett, A., Hammonds, A.S., Ngo, T.-T.B., Misra, S., Murphy, C., Scully, A., Carlson, J.W., Wan, K.H., Laverty, T.R., et al. (2008). Tools for neuroanatomy and neurogenetics in Drosophila. Proc. Natl. Acad. Sci. 105, 9715–9720.

Pfeiffer, B.D., Ngo, T.-T.B., Hibbard, K.L., Murphy, C., Jenett, A., Truman, J.W., and Rubin, G.M. (2010). Refinement of Tools for Targeted Gene Expression in Drosophila. Genetics 186, 735–755.

Prieto-Godino, L.L., Diegelmann, S., and Bate, M. (2012). Embryonic Origin of Olfactory Circuitry in Drosophila: Contact and Activity-Mediated Interactions Pattern Connectivity in the Antennal Lobe. PLoS Biol. 10, e1001400.

R Core Team (2015). R: A Language and Environment for Statistical Computing.

Rauschecker, J.P. (1995). Compensatory plasticity and sensory substitution in the cerebral cortex. Trends Neurosci. 18, 36–43.

Rauschecker, J.P., and Korte, M. (1993). Auditory compensation for early blindness in cat cerebral cortex. J. Neurosci. 13, 4538–4548.

Rees, C.L., Moradi, K., and Ascoli, G.A. (2017). Weighing the Evidence in Peters’ Rule: Does Neuronal Morphology Predict Connectivity? Trends Neurosci. 40, 63–71.

Roberts, A., Conte, D., Hull, M., Merrison-Hort, R., al Azad, A.K., Buhl, E., Borisyuk, R., and Soffe, S.R. (2014). Can simple rules control development of a pioneer vertebrate neuronal network generating behavior? J. Neurosci. 34, 608–621.

Rueden, C.T., Schindelin, J., Hiner, M.C., DeZonia, B.E., Walter, A.E., Arena, E.T., and Eliceiri, K.W. (2017). ImageJ2: ImageJ for the next generation of scientific image data. BMC Bioinformatics 18, 529.

Rutherford, L.C., Nelson, S.B., and Turrigiano, G.G. (1998). BDNF has opposite effects on the quantal amplitude of pyramidal neuron and interneuron excitatory synapses. Neuron 21, 521–530.

Saalfeld, S., Cardona, A., Hartenstein, V., and Tomancak, P. (2009). CATMAID: collaborative annotation toolkit for massive amounts of image data. Bioinformatics 25, 1984–1986.

Sales, E.C., Heckman, E.L., Warren, T.L., and Doe, C.Q. (2019). Regulation of subcellular dendritic synapse specificity by axon guidance cues. Elife 8.

Sanes, J.R., and Yamagata, M. (2009). Many Paths to Synaptic Specificity. Annu. Rev. Cell Dev. Biol.

Schindelin, J., Arganda-Carreras, I., Frise, E., Kaynig, V., Longair, M., Pietzsch, T., Preibisch, S., Rueden, C., Saalfeld, S., Schmid, B., et al. (2012). Fiji: an open-source platform for biological-image analysis. Nat. Methods 9, 676–682.

Schneider-Mizell, C.M., Gerhard, S., Longair, M., Kazimiers, T., Li, F., Zwart, M.F., Champion, A., Midgley, F.M., Fetter, R.D., Saalfeld, S., et al. (2016). Quantitative neuroanatomy for connectomics in Drosophila. Elife 5.

Schulz, D.J., and Lane, B.J. (2017). Homeostatic plasticity of excitability in crustacean central pattern generator networks. Curr. Opin. Neurobiol. 43, 7–14.

Scott, E.K., Reuter, J.E., and Luo, L. (2003). Dendritic development of Drosophila high order visual system neurons is independent of sensory experience. BMC Neurosci. 4, 14.

Shen, K., Fetter, R.D., and Bargmann, C.I. (2004). Synaptic Specificity Is Generated by the Synaptic Guidepost Protein SYG-2 and Its Receptor, SYG-1. Cell 116, 869–881.

Sheng, C., Javed, U., Gibbs, M., Long, C., Yin, J., Qin, B., and Yuan, Q. (2018). Experience-dependent structural plasticity targets dynamic filopodia in regulating dendrite maturation and synaptogenesis. Nat. Commun. 9, 3362.

Sommer, C., Straehle, C., Koethe, U., and Hamprecht, F.A. (2011). ilastik: Interactive Learning and Segmentation Toolkit. In 8th IEEE International Symposium on Biomedical Imaging (ISBI 2011), p.

Sperry, R.W. (1943). Effect of 180 degree rotation of the retinal field on visuomotor coordination. J. Exp. Zool. 92, 263–279.

Sperry, R.W. (1963). CHEMOAFFINITY IN THE ORDERLY GROWTH OF NERVE FIBER PATTERNS AND CONNECTIONS. Proc. Natl. Acad. Sci. 50, 703–710.

Sugie, A., Marchetti, G., and Tavosanis, G. (2018). Structural aspects of plasticity in the nervous system of Drosophila. Neural Dev. 13, 14.

Sürmeli, G., Akay, T., Ippolito, G.C., Tucker, P.W., and Jessell, T.M. (2011). Patterns of spinal sensory-motor connectivity prescribed by a dorsoventral positional template. Cell 147, 653–665.

Sweeney, S.T., Broadie, K., Keane, J., Niemann, H., and O’Kane, C.J. (1995). Targeted expression of tetanus toxin light chain in Drosophila specifically eliminates synaptic transmission and causes behavioral defects. Neuron 14, 341–351.

Takemura, S., Bharioke, A., Lu, Z., Nern, A., Vitaladevuni, S., Rivlin, P.K., Katz, W.T., Olbris, D.J., Plaza, S.M., Winston, P., et al. (2013). A visual motion detection circuit suggested by Drosophila connectomics. Nature 500, 175–181.

Takemura, S., Xu, C.S., Lu, Z., Rivlin, P.K., Parag, T., Olbris, D.J., Plaza, S., Zhao, T., Katz, W.T., Umayam, L., et al. (2015). Synaptic circuits and their variations within different columns in the visual system of Drosophila. Proc. Natl. Acad. Sci. 112, 13711–13716.

Tessier-Lavigne, M., and Goodman, C.S. (1996). The molecular biology of axon guidance. Science 274, 1123–1133.

Tien, N.W., and Kerschensteiner, D. (2018). Homeostatic plasticity in neural development. Neural Dev. 13.

Tripodi, M., Evers, J.F., Mauss, A., Bate, M., and Landgraf, M. (2008). Structural Homeostasis: Compensatory Adjustments of Dendritic Arbor Geometry in Response to Variations of Synaptic Input. PLoS Biol. 6, e260.

Turrigiano, G.G. (2017). The dialectic of Hebb and homeostasis. Philos. Trans. R. Soc. B Biol. Sci. 372.

Turrigiano, G.G., and Nelson, S.B. (2004). Homeostatic plasticity in the developing nervous system. Nat. Rev. Neurosci. 5, 97–107.

del Valle Rodríguez, A., Didiano, D., and Desplan, C. (2012). Power tools for gene expression and clonal analysis in Drosophila. Nat. Methods 9, 47–55.

Venken, K.J.T., Simpson, J.H., and Bellen, H.J. (2011). Genetic Manipulation of Genes and Cells in the Nervous System of the Fruit Fly. Neuron 72, 202–230.

Vogelstein, J.T., Park, Y., Ohyama, T., Kerr, R.A., Truman, J.W., Priebe, C.E., and Zlatic, M. (2014). Discovery of brainwide neural-behavioral maps via multiscale unsupervised structure learning. Science 344, 386–392.

Vonhoff, F., and Keshishian, H. (2017). In Vivo Calcium Signaling during Synaptic Refinement at the Drosophila Neuromuscular Junction. J. Neurosci. 37, 5511–5526.

Ward, A., Hong, W., Favaloro, V., and Luo, L. (2015). Toll receptors instruct axon and dendrite targeting and participate in synaptic partner matching in a drosophila olfactory circuit. Neuron 85, 1013–1028.

White, J.G., Southgate, E., Thomson, J.N., and Brenner, S. (1986). The structure of the nervous system of the nematode Caenorhabditis elegans. Philos. Trans. R. Soc. Lond. B. Biol. Sci. 314, 1–340.

Wickham, H. (2009). ggplot2: Elegant Graphics for Data Analysis (New York: Springer-Verlag).

Wolfram, V., and Baines, R.A. (2013). Blurring the boundaries: developmental and activity-dependent determinants of neural circuits. Trends Neurosci. 36, 610–619.

Yogev, S., and Shen, K. (2014). Cellular and Molecular Mechanisms of Synaptic Specificity. Annu. Rev. Cell Dev. Biol 30, 417–437.

Yuan, Q., Xiang, Y., Yan, Z., Han, C., Jan, L.Y., and Jan, Y.N. (2011). Light-induced structural and functional plasticity in Drosophila larval visual system. Science 333, 1458–1462.

Zheng, Z., Lauritzen, J.S., Perlman, E., Robinson, C.G., Nichols, M., Milkie, D., Torrens, O., Price, J., Fisher, C.B., Sharifi, N., et al. (2018). A Complete Electron Microscopy Volume of the Brain of Adult Drosophila melanogaster. Cell 174, 730–743.e22.

Zhu, H., Hummel, T., Clemens, J.C., Berdnik, D., Zipursky, S.L., and Luo, L. (2006). Dendritic patterning by Dscam and synaptic partner matching in the Drosophila antennal lobe. Nat. Neurosci. 9, 349–355.

Zlatic, M., Landgraf, M., and Bate, M. (2003). Genetic Specification of Axonal Arbors: atonal Regulates robo3 to Position Terminal Branches in the Drosophila Nervous System. Neuron 37, 41–51.

Zlatic, M., Li, F., Strigini, M., Grueber, W., and Bate, M. (2009). Positional Cues in the Drosophila Nerve Cord: Semaphorins Pattern the Dorso-Ventral Axis. PLoS Biol. 7, e1000135.

